# Deletion of RFX6, a Diabetes-Associated Gene, Impairs iPSC-Derived Islet Organoid Development and Survival, With No Impact on the Generation of PDX1+/NKX6.1+ Progenitors

**DOI:** 10.1101/2024.03.06.583663

**Authors:** Noura Aldous, Ahmed K. Elsayed, Bushra Memon, Sadaf Ijaz, Sikander Hayat, Essam M. Abdelalim

**Affiliations:** College of Health and Life Sciences, Hamad Bin Khalifa University (HBKU), Qatar Foundation, Education City, Doha, Qatar; Diabetes Research Center, Qatar Biomedical Research Institute (QBRI), Hamad Bin Khalifa University (HBKU), Qatar Foundation (QF), PO Box 34110, Doha, Qatar; Pluripotent Stem Cell Disease Modeling Lab, Translational Medicine Division, Research Branch, Sidra Medicine, P.O. Box 26999, Doha, Qatar; Stem Cell Core, Qatar Biomedical Research Institute (QBRI), Hamad Bin Khalifa University (HBKU), Qatar Foundation (QF), PO Box 34110, Doha, Qatar; Department of Medicine 2, Medical Faculty RWTH Aachen University, Germany

**Keywords:** transcription factors, endocrine specification, pancreatic progenitors, islet organoids, pancreatic hypoplasia, diabetes

## Abstract

RFX6 is essential for pancreatic development and insulin secretion, while its role in diabetes pathogenesis is unclear. Here, RFX6 expression was detected in PDX1+ cells in the hESC-derived posterior foregut (PF). However, in the pancreatic progenitors (PPs), RFX6 did not co-localize with PDX1 and NKX6.1, but instead with NEUROG3, NKX2.2, and islet hormones in the endocrine progenitor (EPs) and islets. Single-cell analysis revealed high RFX6 expression in endocrine clusters across various hESC-derived pancreatic differentiation stages. Upon differentiating iPSCs lacking RFX6 into pancreatic islets, a significant decrease in PDX1 expression at the PF stage was observed, although it did not affect PPs co-expressing PDX1 and NKX6.1. RNA sequencing showed the downregulation of essential genes involved in pancreatic endocrine differentiation, insulin secretion, and ion transport due to RFX6 deficiency. Furthermore, RFX6 deficiency resulted in the formation of smaller islet organoids due to increased cellular apoptosis, linked to reduced Catalase (CAT) expression, implying a protective role for RFX6. Overexpression of RFX6 reversed defective phenotypes in PPs and EPs. These findings suggest that pancreatic hypoplasia and reduced islet cell formation associated with RFX6 mutations are not due to alterations in PDX1+/NKX6.1+ PPs but instead result from cellular apoptosis and downregulation of pancreatic endocrine genes.

## Introduction

RFX6, a transcription factor, is highly expressed in the pancreas and plays a crucial role in the development of islet cells. Homozygous mutations in RFX6 lead to Mitchell-Riley syndrome (MRS), an autosomal recessive disorder characterized by severe neonatal diabetes, hypoplastic or annular pancreas, and intestinal atresia [1–5]. It has been suggested that the diabetes observed in individuals with homozygous RFX6 mutations is attributed to the overall impairment of pancreatic islet development and function, including a reduction in insulin production by β-cells [1, 6]. In contrast, heterozygous RFX6 mutations can cause mild maturity-onset diabetes of the young (MODY) with reduced penetrance [7–9]. In these patients with RFX6-MODY, defective insulin secretion is attributed to the failure of islet β-cells to properly secrete insulin, despite the normal development of the islets [4, 10]. Thus, it appears that RFX6 plays a dual role in controlling both pancreatic islet development and insulin production, but through distinct mechanisms. Studies involving RFX6 knockdown in human EndoC-βH1 cells have demonstrated changes in insulin mRNA levels [4]. Genome-wide association studies (GWAS) identified RFX6 variants associated with type 2 diabetes (T2D) [11]. The integration of genomic, epigenomic, and transcriptomic data from human islets of T2D patients has identified genetic loci that show specific enrichment of RFX-binding motifs [12].

Studies in mice have revealed that Rfx6 expression initiates in the gut endoderm, undergoes a developmental transition, and eventually becomes restricted to the endocrine lineage within the pancreas, persisting in adult islet cells. The significance of Rfx6 in islet cell development is evident across various species, including zebrafish [13], Xenopus [6], mice [1], and humans [1, 2, 6, 14]. In mice with Rfx6 gene deficiency, all endocrine cells, including β-cells, are absent, except for polypeptide-secreting (PP) cells, resulting in diabetes and early postnatal death in mice, which limits the exploration of its role in β-cell function and insulin production [1, 6]. Moreover, in adult β-cells, the loss of Rfx6 results in glucose intolerance, impaired glucose sensing, and defective insulin secretion [10]. Previous research highlighted the expression of Rfx6 in GIP-positive enteroendocrine K-cells and its role in regulating Gip promoter activity [15]. Furthermore, a recent study demonstrated that the loss of Rfx6 function in *ex vivo* mouse intestinal organoids led to a reduction in enteroendocrine cells (EECs) [16].

The existence of various forms of diabetes among individuals harbouring RFX6 mutations [1] prompts inquiries about the necessity of RFX6 in the development and function of pancreatic islets. This potential function warrants thorough exploration and could potentially identify RFX6 as a promising therapeutic target for diabetes. While the previous data indicate that RFX6 plays a crucial role in the development of pancreatic islets and contributes to maintaining physiological glucose homeostasis, the changes in the expression and/or function of RFX6 during pancreatic islet development and its precise involvement in diabetes pathogenesis remain unclear. In this study, we established induced pluripotent stem cell (iPSC) lines with RFX6 loss of function to unravel its significance in the development of human islet cells, employing a CRISPR/Cas9 technique and a stepwise differentiation protocol. RFX6 loss exhibited no significant impairments in the capacity of iPSCs to generate pancreatic progenitors (PPs), co-expressing PDX1 and NKX6.1. However, the lack of RFX6 was associated with significant decrease in the expression of early pancreatic endocrine specification genes and an increased cell death in iPSC-derived pancreatic islet organoids with significant reduction in the Catalase (CAT) enzyme. Our findings indicate that RFX6 plays a vital role in regulating specific genes related to pancreatic endocrine development and is essential for islet organoid survival during the late stages of iPSC-derived islet development.

## Results

### Temporal and cell-specific expression of RFX6 during pancreatic islet differentiation

To characterize the expression of RFX6 throughout the pancreatic islet differentiation process, we differentiated hESC-H9 cells into pancreatic islets and examined the expression of RFX6 at various stages during the pancreatic differentiation (Supplementary Fig. 1A). As of now, there is no available specific antibody for studying the RFX6 protein expression using immunostaining. Instead, we employed a recently developed modified hESC-H9 line, known as *RFX6^HA/HA^* hESC-H9, where a triplicated HA epitope is inserted into the 3’ end of the RFX6 gene to facilitate the investigation of RFX6 protein expression [17]. Using immunostaining, RFX6 expression was not detected during the definitive endoderm (DE) and primitive gut tube (PGT) stages (Supplementary Fig. 1B). However, robust RFX6 expression was detected in the posterior foregut (PF), where it largely co-localized with PDX1 (Fig. 1A). Surprisingly, RFX6 expression continued at the pancreatic progenitor (PP) stage but did not co-express with the hallmark transcriptional factors PDX1 and NKX6.1 (Fig. 1A). During the endocrine progenitor (EP) stage, RFX6 was co-localized with NEUROG3, the master initiator of the pancreatic endocrine lineage, and also co-expressed with another endocrine marker, NKX2.2 (Fig. 1A). In the islet cell stage, RFX6 showed co-expression with insulin (INS), glucagon (GCG), and somatostatin (SST) (Fig. 1A). These results were further validated by flow cytometry quantification, showed that high RFX6 protein expression started at the PF stage, continued consistently with a peak observed at the EP stage (Fig. 1B). These results suggest that RFX6 may be not essential for the PDX1+/NKX6.1+ PPs.

**Figure 1.**
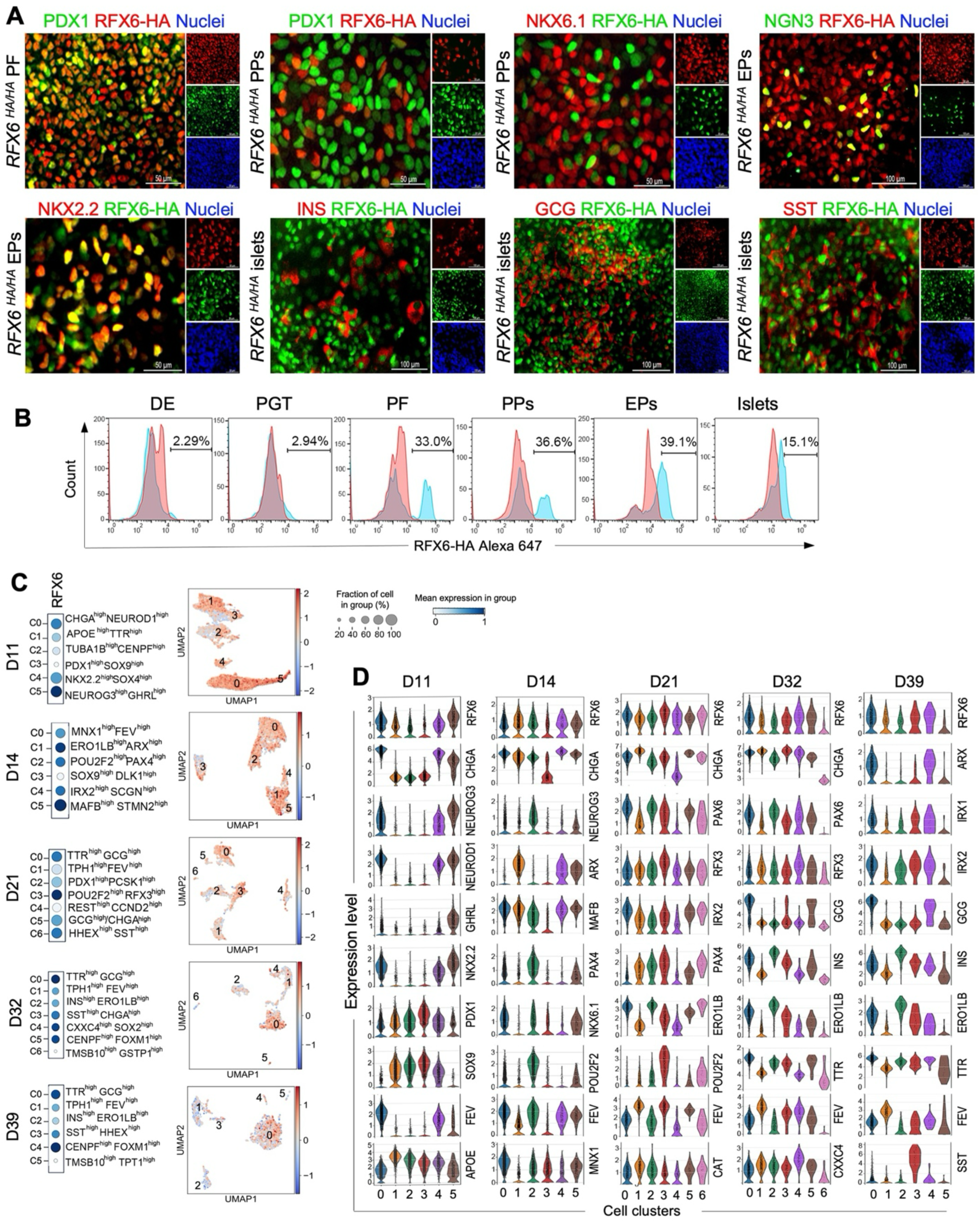
Timeline expression and single cell analysis of RFX6 throughout the differentiation of hESCs into various stages of pancreatic development. **(A)** Immunostaining showing the expression of RFX6 during differentiation of hESC-H9 into pancreatic islets. **(B)** Flow cytometric quantification of RFX6 expression during different stages of differentiation. **(C)** Dot plots and feature plots illustrating RFX6 expression across distinct cell clusters. Each dot’s color and size correspond to the expression level and the percentage of cells expressing RFX6 gene. **(D)** The violin plots illustrate the expression distributions of key genes across various clusters at distinct stages of hESC differentiation into pancreatic islets: day 11 (D11), day 14 (D14), day 21 (D21), day 32 (D32), and day 39 (D39). DE: definitive endoderm, PGT: primitive gut tube, PF: posterior foregut, PPs: pancreatic progenitors, EPs: endocrine progenitors. Scale bars = 100 µm.

To delve deeper into identifying specific cell populations expressing RFX6, we re-analyzed the recently published single-cell (scRNA-seq) datasets of PPs at day 11 (D11), EPs at day 14 (D14), immature islets at day 21 (D21), and maturing islets at days 32 and 39 (D32 and D39), derived from hESCs [18]. We used unsupervised clustering to create 2D visualizations using Uniform Manifold Approximation and Projection (UMAP) plots. Through analyzing RNA expression patterns, we identified multiple cell populations at each stage of differentiation (Fig. 1C, D; Supplementary Fig. 2). At D11, we identified six distinct cell clusters, with three of them showed high expression of pancreatic endocrine markers. Interestingly, RFX6 was mainly expressed in these three endocrine clusters with the highest in C5, distinguished by NEUROG3^high^/GHRL^high^, which also expressed high levels of other endocrine markers such as PAX4, INSM1, KCNK17, NKX2.2, and SOX4. Furthermore, moderate levels of RFX6 expression were observed in other two endocrine clusters (C0 and C4) characterized by CHGA^high^/NEUROD1^high^ and NKX2.2^high^/SOX4^high^, respectively. A small number of cells expressed RFX6 in C1 (APOE^high^/TTR^high^) and C2 (proliferation cluster; TUBA1B^high^/CENPF^high^). However, almost no expression was detected in C3, identified by PDX1^high^/SOX9^high^ (Fig. 1C, D; Supplementary Fig. 2). These findings strongly indicate that RFX6 expression in PPs is confined to the endocrine cell populations.

At D14, we also identified six clusters, with the greatest expression of RFX6 was seen within the endocrine cluster C5, marked by MAFB^high^/STMN2^high^. This clusters also showcased elevated levels of essential endocrine markers like INS, GCG, and SLC30A8 (Fig. 1C, D; Supplementary Fig. 2). Also, high RFX6 expression was seen in another endocrine cluster, C1 (ERO1LB^high^/ARX^high^). In addition, a moderate RFX6 level was detected in C2 (POUF2F2^high^/PAX4^high^), C0 (MNX1^high^/ FEV^high^), and C4 (IRX2^high^/SCGN^high^). The lowest RFX6 expression seen in C3 (pancreatic progenitor cluster; SOX9^high^/DLK1^high^), which also expressed high levels of PDX1, HNF1B, GATA4, TCF7L2, and CCND2. At D21, the highest expression of RFX6 was seen in C3 (POU2F2^high^/RFX3^high^), while a moderate expression was seen in C0 (TTR^high^/GCG^high^), C5 (GCG^high^/CHGA^high^), and C6 (δ-cell cluster; HHEX^high^/SST^high^). Reduced expression was observed in C1 (TPH1^high^/FEV^high^), previously identified as a specific enterochromaffin progenitor population [19], and in C4 (proliferation cluster; REST^high^/ CCND2^high^) (Fig. 1C, D; Supplementary Fig. S2).

At D32, the most significant RFX6 expression was seen in C0 (TTR^high^/GCG^high^), which also expressed high levels of CHGA, IRX2, and ARX, suggesting an α-cell fate. Moderate RFX6 expression levels were seen in C4 (CXXC4^high^/SOX2^high^) and C5 (proliferation cluster; CENPF^high^/FOXM1^high^), while comparatively lower expression was seen in C1 (TPH1^high^/FEV^high^) and C2 (INS^high^/ERO1LB^high^). ERO1LB is known as a gene specifically associated with pancreatic β-cells.[20] No expression was observed in C6 (TMSB10^high^/GSTP1^high^) (Fig. 1C, D; Supplementary Fig. 2). At D39, the highest expression of RFX6 was seen in C4 (proliferation cluster; CENPF^high^/FOXM1^high^), with moderate expression was noted in C0 (TTR^high^/GCG^high^). Lower expression was seen in C3 (SST^high^/HHEX^high^), C1 (TPH1^high^/FEV^high^), and C2 (INS^high^/ERO1LB^high^), while no expression was seen in C5 (TMSB10^high^/TPT1^high^) (Fig. 1C, D; Supplementary Fig. 2). Taken together, these findings indicate that RFX6 is mainly expressed in pancreatic endocrine cell populations across various stages and is not expressed in PDX1+ cell populations within pancreatic progenitors.

### Depletion of RFX6 diminishes PDX1 expression in the posterior foregut and does not affect PDX1+/NKX6.1+ pancreatic progenitors

To explore RFX6’s contribution to human pancreatic development and the generation of pancreatic islet cells, biallelic RFX6 mutant hiPSC lines (referred to as RFX6 KO-iPSCs) were established using the CRISPR/Cas9 system. Mutations were introduced into the wild type (WT) iPSC line (referred to as WT-iPSCs) generated in our laboratory [21] using a sgRNA that targeted exon 2 of the RFX6 gene. The mutations were validated through Sanger sequencing and were anticipated to induce a frameshift, resulting in the formation of premature stop codons preventing RFX6 protein translation (Fig. 2A). The absence of RFX6 protein expression was confirmed in PPs derived from RFX6 KO cell lines compared to WT controls using Western blot analysis (Fig. 2B). All iPSC lines expressed pluripotency markers OCT4, NANOG, SOX2, SSEA4, TRA-1-60, TRA-81, C-MYC, KLF4, REX1, DPPA4, and TERT (Supplementary Fig. 3A, B). Moreover, they have been verified to maintain a normal karyotype consistent with the parental line and are free from mycoplasma (Supplementary Fig. 3C, D)

**Figure 2.**
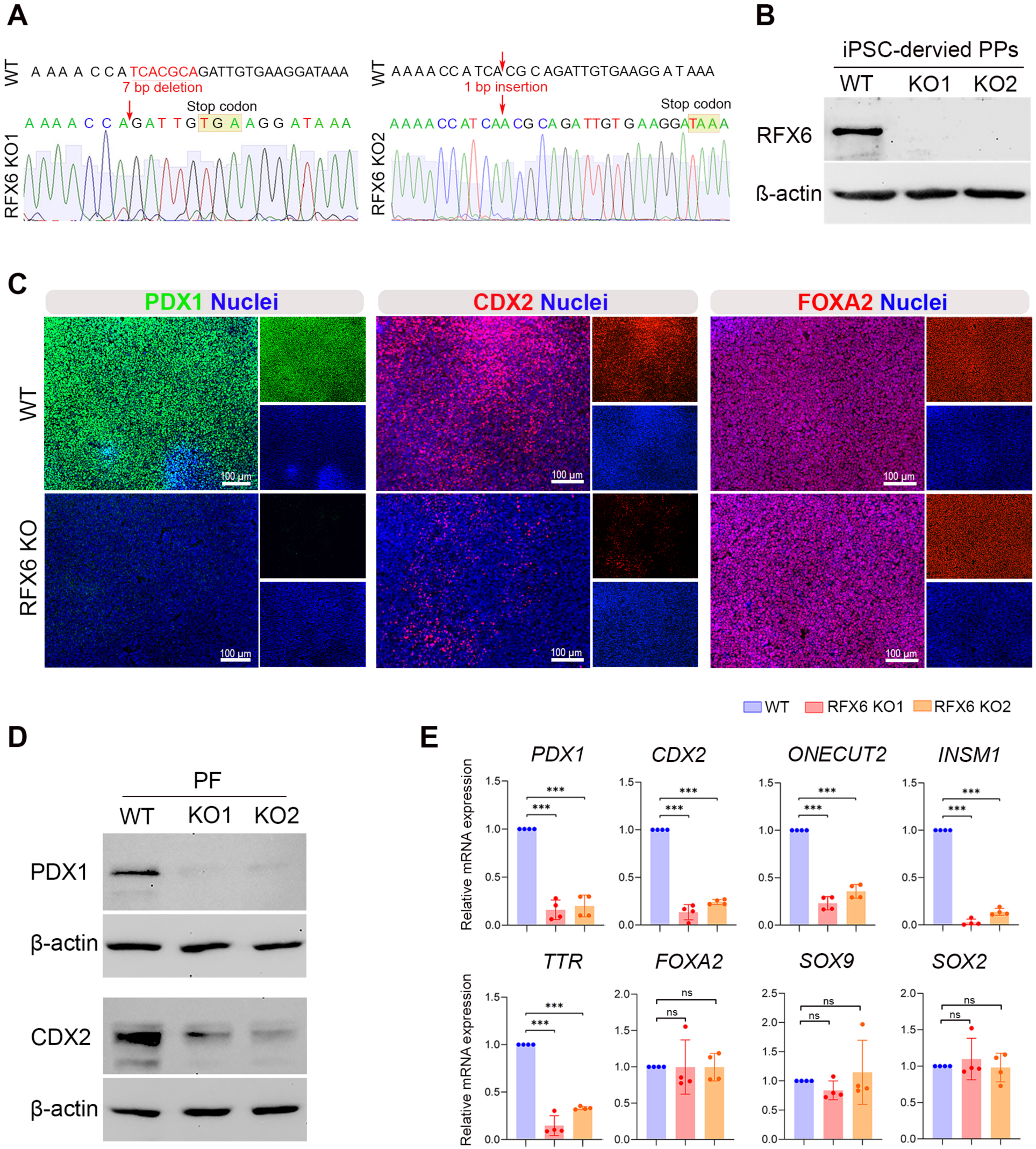
Loss of RFX6 reduces PDX1 and CDX2 expression in iPSC-derived posterior foregut. **(A)** DNA sequence confirmation of frameshift mutations in isogenic KO iPSC clones compared to WT iPSCs. **(B)** Western blot analysis confirming the absence of RFX6 protein in pancreatic progenitors (PPs) derived from RFX6 KO iPSC lines. **(C)** Immunofluorescence images showing the expression of PDX1, CDX2, and FOXA2 in PPs derived from WT-iPSCs and RFX6 KO iPSCs. **(D)** Western blot analysis showing the expression of PDX1 and CDX2 in RFX6 KO posterior foregut (PF) compared to WT-PF. **(E)** RT-qPCR analysis showing the mRNA expression of PF markers*, PDX1*, *CDX2*, *ONECUT2*, *INSM1, TTR, FOXA2, SOX9,* and *SOX2* in RFX6 KO PF relative to wild type (WT) controls (n=4). The data are presented as mean ±SD. \**p* < 0.05, \*\**p* < 0.01, \*\*\**p* < 0.001. Scale bar = 100 µm.

Next, we examined the effect of RFX6 loss on pancreatic differentiation stages that express RFX6. Immunostaining and Western blotting showed that the lack of RFX6 resulted in significant reduction in PDX1 and CDX2 protein expressions, whereas FOXA2 expression remained unchanged in PFs (Fig. 2C, D). Also, RT-qPCR analysis showed significant downregulation in the mRNA expression of *PDX1, CDX2, ONECUT2, INSM1,* and *TTR* in RFX6 KO-PFs compared to WT-PFs (Fig. 2E). On the other hand, the absence of RFX6 had no significant impact on the expression of *FOXA2, SOX9,* and *SOX2* (Fig. 2E). Interestingly, despite the dramatic reduction in PDX1 expression during the PF stage, RFX6-deficient iPSCs were able to produce PPs that exhibited the co-expression of PDX1 and NKX6.1, similar to the WT controls (Fig. 3A-D). Furthermore, there were no significant changes in the expression of other PP markers, including SOX9 and FOXA2, as evidenced by Western blotting and RT-qPCR (Figure 3C, D). These results suggest that RFX6 does not play a significant role in the formation of PDX1^+^/NKX6.1^+^ cells during the PP stage.

**Figure 3.**
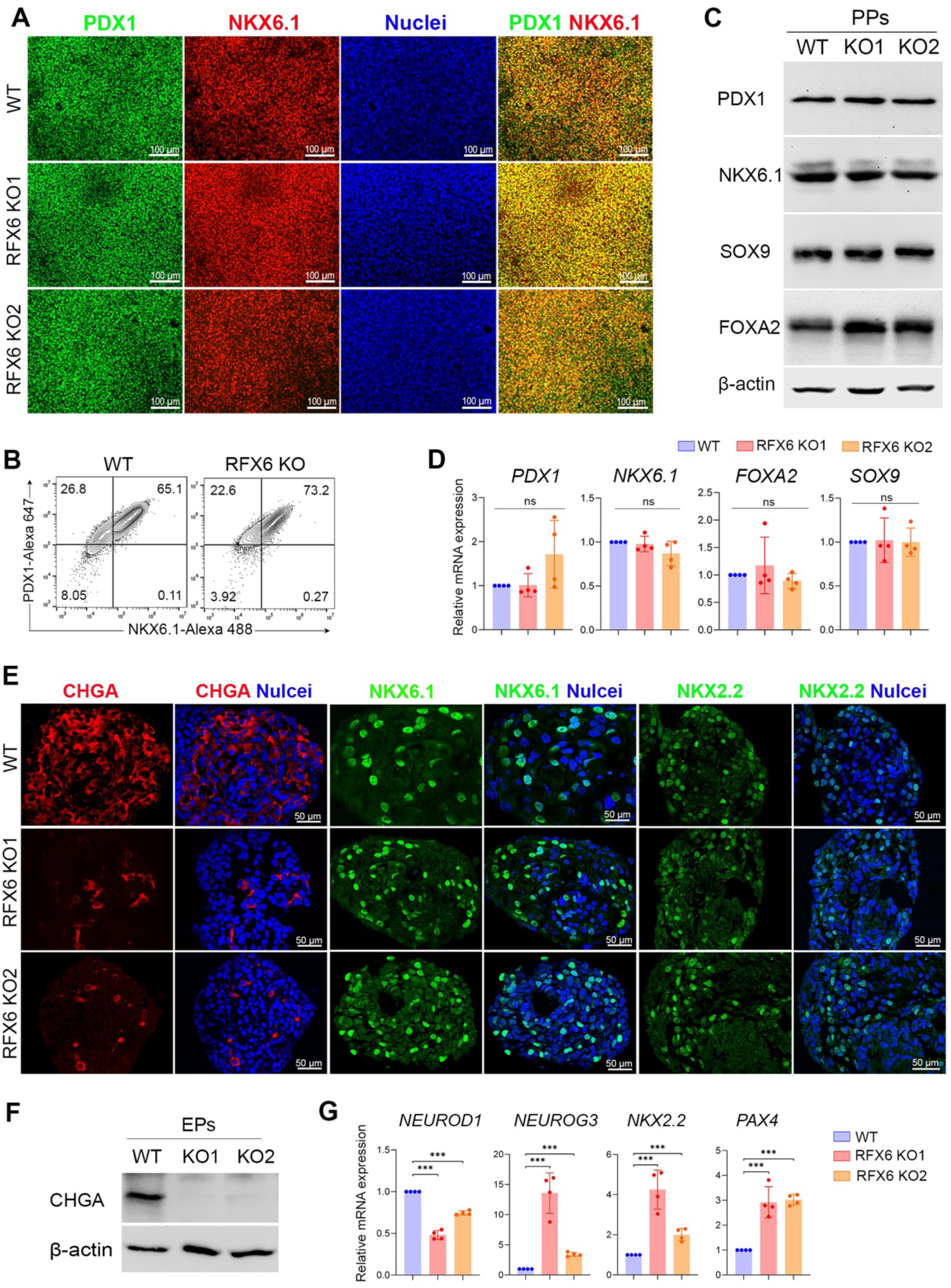
Impact of RFX6 depletion on the expression of crucial pancreatic progenitor and endocrine progenitor markers. Immunofluorescence staining **(A)** and flow cytometry analysis **(B)** showing the co-expression of PDX1 and NKX6.1 in PPs derived from WT-iPSCs and RFX6 KO iPSCs. **(C)** Western blot analysis showing the protein expression of PDX1, NKX6.1, SOX9, and FOXA2 in RFX6 KO PPs compared to WT-PPs. **(D)** RT-qPCR analysis showing the mRNA expression of PP markers*, PDX1, NKX6.1, FOXA2,* and *SOX9* in RFX6 KO PPs relative to WT-PPs (n=4). **(E)** Immunofluorescence staining showing the expression of CHGA, NKX6.1, and NKX2.2 in EPs derived from WT-iPSCs and RFX6 KO iPSCs. **(F)** Western blot analysis showing the expression of CHGA in RFX6 KO EPs compared to WT-EPs. (**G)** RT-qPCR analysis showing the mRNA expression of EP markers*, NEUROD1, NEUROG3, NKX2.2,* and *PAX4* in RFX6 KO EPs relative to WT-EPs (n=4). The data are presented as mean ±SD. \**p* < 0.05, \*\**p* < 0.01, \*\*\**p* < 0.001. Scale bars = 50, 100 µm.

At the EP stage, our immunostaining and Western blotting results showed a dramatic reduction of the pan endocrine marker, CHGA (Fig. 3E, F); however, there was no significant change in the expression of NKX6.1 and NKX2.2 by immunostaining (Fig. 3E). RT-qPCR analysis showed significant reduction in the expression of *NEUROD1*, while the expression of *NEUROG3, NKX2.2,* and *PAX4* were significantly increased in EPs lacking RFX6 compared to WT controls (Fig. 3G).

### Deletion of RFX6 leads to large-scale transcriptomic alterations associated with pancreatic endocrine specification in pancreatic progenitors and endocrine progenitors

For comprehensive understanding of the transcriptomic changes between RFX6 KO and WT cells, RNA sequencing (RNA-seq) was performed on PPs and EPs. Our transcriptome analysis on iPSC-derived PPs, detected 393 differentially expressed genes (DEGs) significantly affected by RFX6 deletion. Among these DEGs, 224 genes were significantly downregulated (Log2 FC < −1.0, *p* < 0.05), while 169 genes were significantly upregulated (Log2 FC > 1.0, *p* < 0.05) in RFX6 KO-PPs compared to WT-PPs (Fig. 4A; Supplementary Fig. 4A). At the EP stage, we identified 334 DEGs significantly impacted by the deletion of RFX6, with 215 of these genes were significantly downregulated (Log2 FC < −1.0, *p* < 0.05), and 119 genes were significantly upregulated (Log2 FC > 1.0, *p* < 0.05) in RFX6 KO-EPs compared to WT-EPs (Fig. 4A; Supplementary Fig. 4A). Interestingly, 160 of the downregulated DEGs, comprising 57.3%, were found in both PPs and EPs (Fig. 4B), with most of these genes known to be associated with pancreatic endocrine development. The Gene Ontology (GO) of the downregulated DEGs in PPs and EPs displayed enriched genes linked to pancreatic endocrine development, insulin secretion regulation, and regulation of ion transmembrane transport (Fig. 4C; Supplementary Fig. 4C), whereas the upregulated DEGs showed GO enrichment in lipid metabolism and nervous system development (data not shown). At PP stage, the RT-qPCR validation analysis confirmed a significant decrease in the expression of endocrine genes, including *ARX, PAX6, CHGA, IRX1, IRX2, INS, GCG, SST, MAF1B, ERO1B (ERO1LB), NEUROD1, PCSK1, CRYBA2, SCGN, PTPRN, PRPRN2, FEV,* and *LMX1B* in RFX6 KO-PPs compared to WT-PPs (Fig. 4D; Table 1). Furthermore, at the EP stage, the RT-qPCR revealed significant decrease in the expression of endocrine genes, including *ARX, PAX6, ISL1, IRX2, INS, GCG, SST, NEUROD1, PCSK1, SCGN, ERO1B, MAFB, SIX3, KCTD12,* and *LMX1B* in RFX6 KO-EPs compared to WT-EPs (Fig. 4E; Table 1).

**Figure 4.**
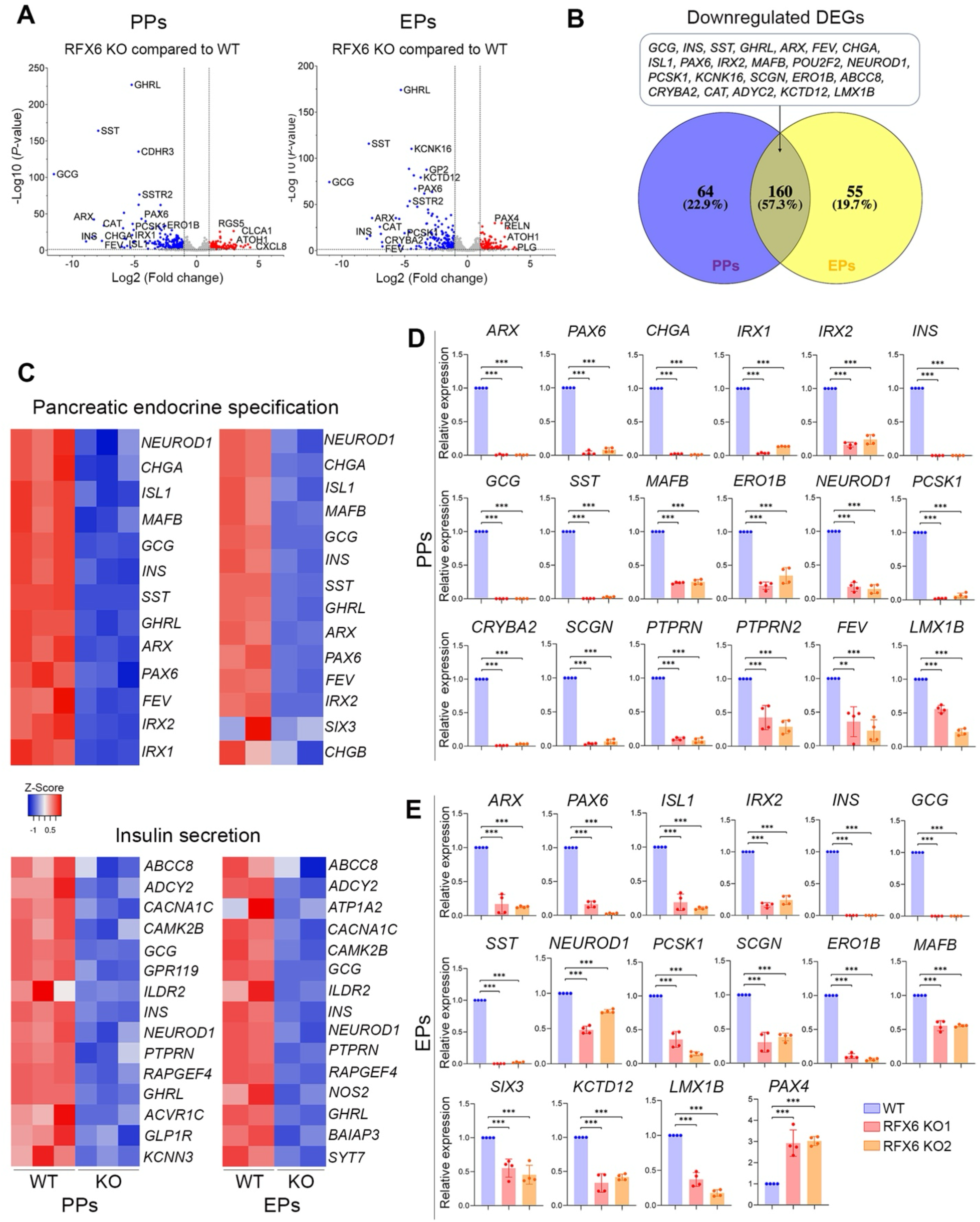
Impact of RFX6 loss on transcriptomic profiles of iPSC-derived pancreatic progenitors (PPs) and endocrine progenitors (EPs). Bulk RNA-seq analysis was performed on pancreatic progenitors (PPs) (n=3) and endocrine progenitors (EPs) (n=2) derived from RFX6 KO iPSCs and WT-iPSCs. **(A)** Volcano plots display the differentially expressed genes (DEGs) in RFX6 KO PPs and RFX6 KO EPs compared to their WT controls. Downregulated genes are represented by blue dots, while upregulated genes are depicted by red dots. **(B)** Venn diagram illustrating the intersection of downregulated DEGs in RFX6 KO PPs and RFX6 KO EPs. Note that most of those DEGs are endocrine pancreatic genes. **(C)** Heatmap of *Z* score value of pancreatic endocrine and insulin secretion genes downregulated in RFX6 KO PPs, and RFX6 KO EPs compared to WT-PPs and WT-EPs, respectively. RT-qPCR validation of the DEGs in PPs **(D)** and EPs **(E)** derived from two different KO iPSC lines (n=4). The data are presented as mean ±SD. \**p* < 0.05, \*\**p* < 0.01, \*\*\**p* < 0.001.

**Table 1.**
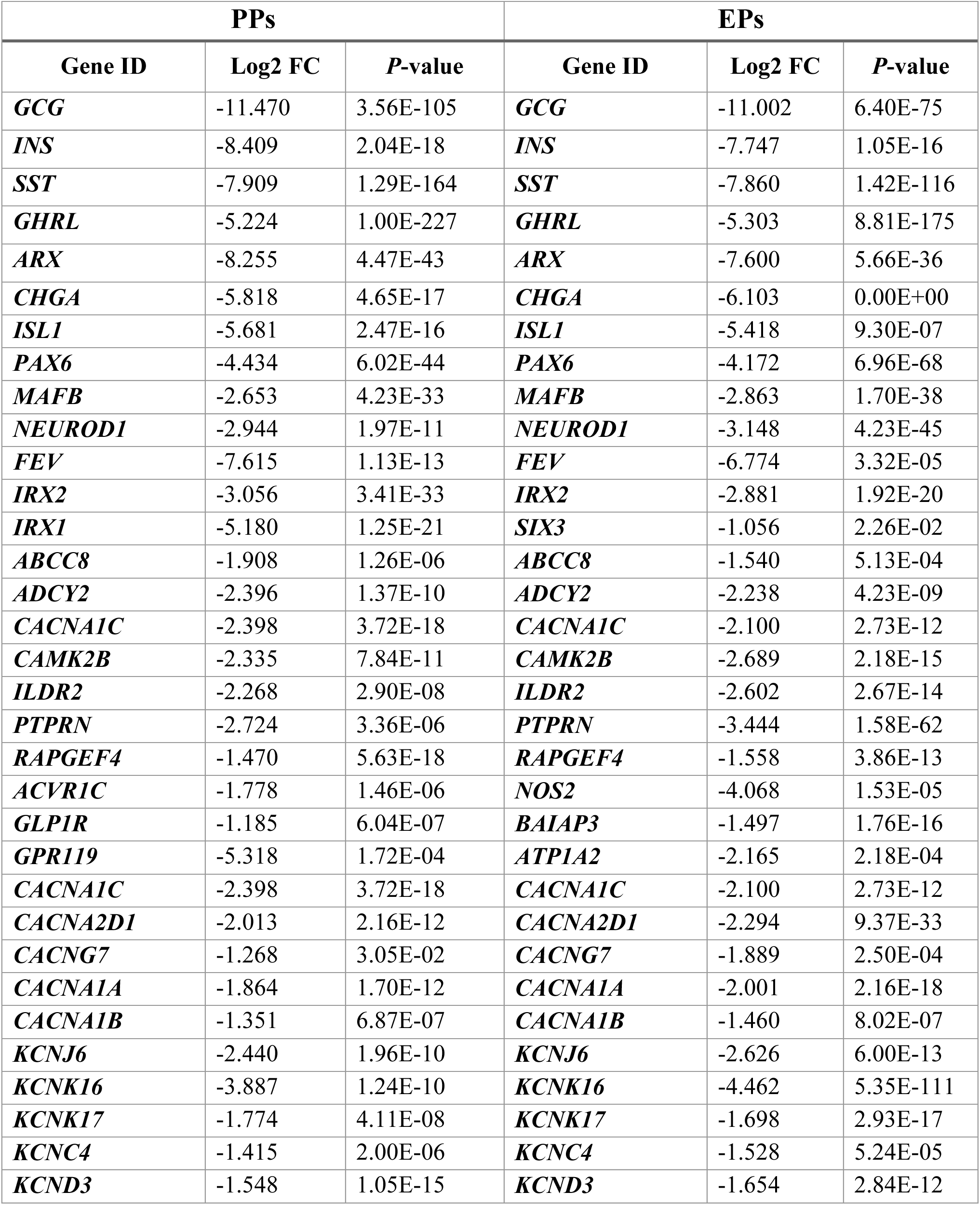
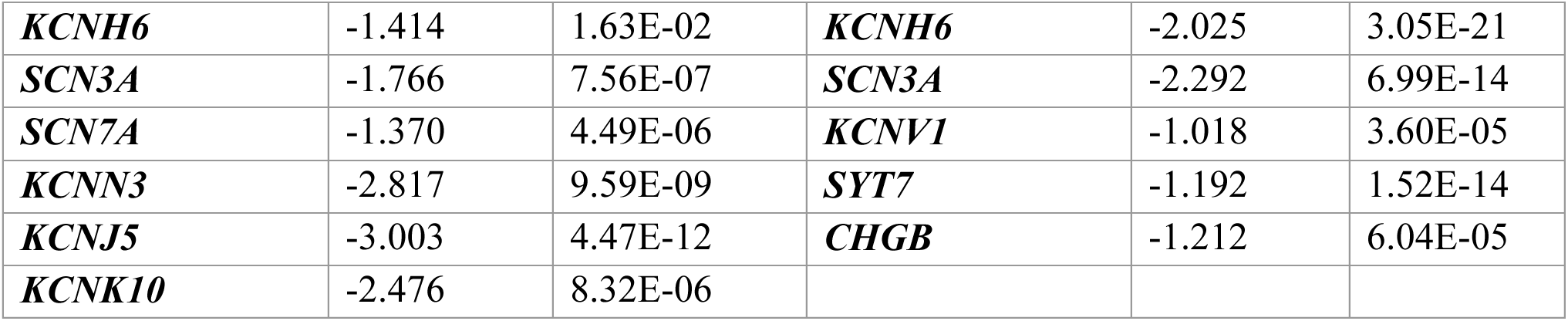
Key downregulated differentially expressed genes (DEGs) associated with pancreatic endocrine development and function in PPs and EPs lacking RFX6 (Log2 FC < –1, *P*-value <0.05).

### RFX6 loss correlates with the generation of smaller pancreatic islet organoids

To enhance islet differentiation after stage 4, cells were cultured in suspension to form organoids, with an equal number of WT-PP and KO-PP cells used. Although during the first two days of stage 5, no notable difference between WT and KO organoids was observed, a significant variation in organoids size became evident as differentiation progressed. Islet organoids derived from RFX6 KO iPSCs showed smaller sizes and irregular shapes compared to those derived from WT-iPSCs during stages 5 and 6 (Fig. 5A).

**Figure 5.**
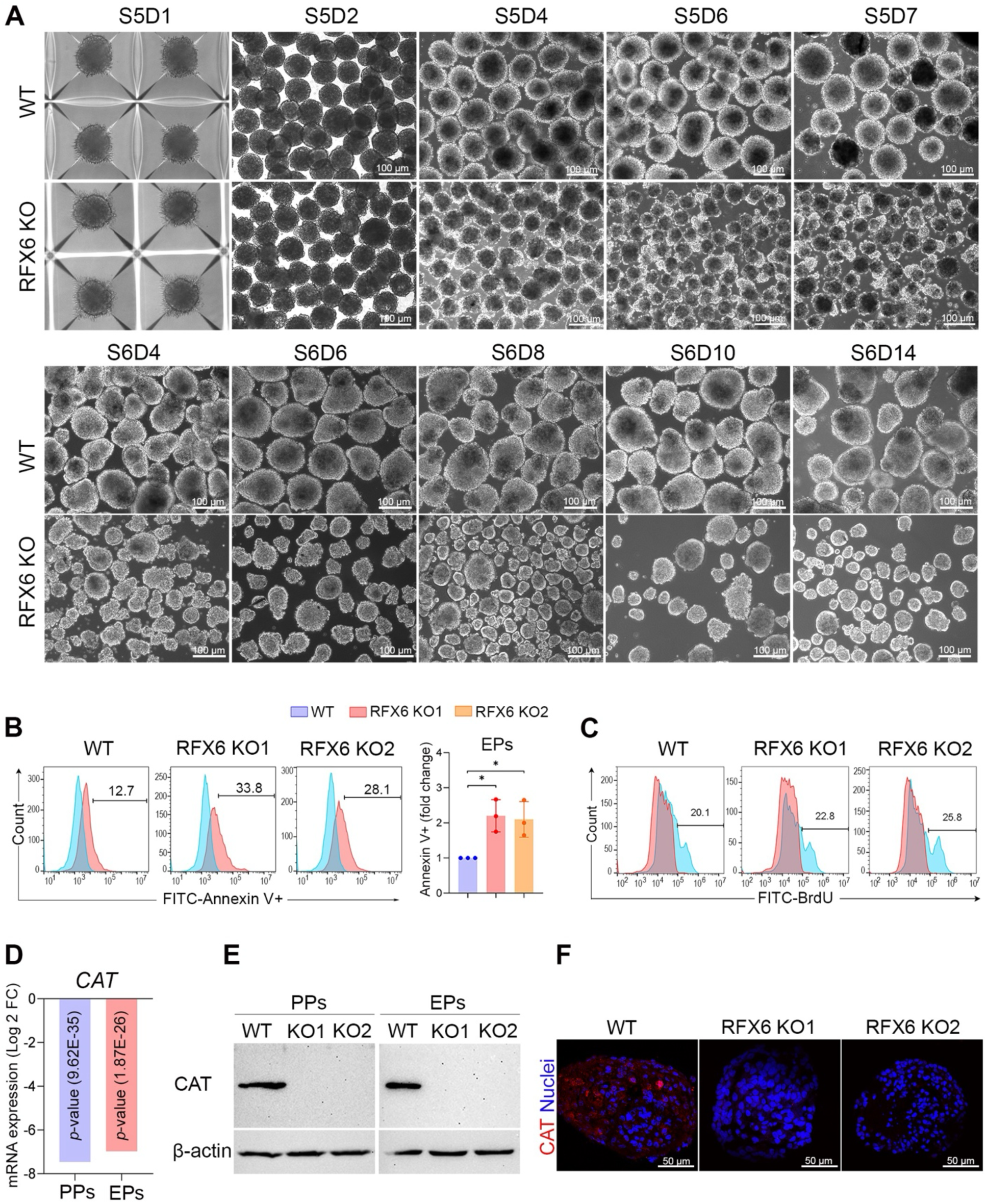
Influence of RFX6 deletion on pancreatic islet organoid formation and cell viability. (**A**) Comparative morphological analysis of pancreatic islet organoids derived from two RFX6 KO iPSC lines versus WT-iPSCs during differentiation stages 5 and 6 (n=3). **(B)** Representative flow cytometry analysis and quantification of apoptosis (Annexin V+ cells) on day 3 of stage 5 of differentiation indicates a significant increase in apoptosis in RFX6 KO EPs in comparison to WT-EPs (n=3). (**C**) Flow cytometry analysis of BrdU incorporation reveals a slight increase in cell proliferation (BrdU+ cells) in endocrine progenitors (EPs) derived from RFX6 KO iPSC lines compared to those derived from WT-iPSCs. **(D)** Log2 fold change in the expression of *CAT* mRNA in RFX6 KO PPs and RFX6 KO EPs compared to WT controls, based on RNA-seq data analysis. **(E)** Western blot analysis showing the absence of CAT protein in RFX6 KO PPs and RFX6 KO EPs compared to WT controls. **(F)** Immunofluorescence images showing the lack of CAT expression in RFX6 KO EPs compared to WT-EPs. The data are presented as mean ±SD. \**p* < 0.05, \*\**p* < 0.01, \*\*\**p* < 0.001. Scale bars = 50 µm, 100 µm.

To investigate whether the dramatic reduction in islet organoid size is attributed to either cell death or inhibition of cell proliferation, we conducted apoptosis and proliferation assays during stage 5 of differentiation. Cell viability was assessed using Annexin V staining. Flow cytometry analysis demonstrated a significant increase in the proportion of Annexin V+ cells in RFX6 KO-EPs compared to WT-EPs (Fig. 5B). However, quantification of BrdU incorporation via flow cytometry revealed no significant difference in the proliferation rate between WT and KO cells (Fig. 5C). This suggests that the reduced size of islet organoids associated with RFX6 loss is mainly due to increased cell death.

In order to elucidate the mechanism underlying the increased cell death, we analyzed the top DEGs identified from our RNA-seq data. Interestingly, we observed a significant downregulation of the antioxidant enzyme *CAT*, which is known to protect cells against oxidative stress [22], in both RFX6 KO-PPs (Log2 FC = -7.459; *p*-value = 9.62E-35) and RFX6 KO-EPs (Log2 FC = - 6.978; *p*-value = 1.87E-26) compared to WT controls (Fig. 5D). This finding was validated at the protein level through Western blot and immunostaining analyses, revealing an almost complete absence of CAT expression in both RFX6 KO-PPs and RFX6 KO-EPs compared to their respective WT controls (Fig. 5E, F). Validating the role of the CAT enzyme in promoting cell survival, we employed the STRING tool that predicts functional interactions of proteins [23]. The analysis revealed that CAT predominantly interacts with proteins involved in safeguarding cells against oxidative stress, such as superoxide dismutase proteins (SOD1, SOD3, and SOD2-2) (Supplementary Fig. 4D).

### RFX6 loss hinder the development of pancreatic islet cells

Subsequent differentiation into pancreatic islets demonstrated a lack of expression for INS, Proinsulin (PROINS), GCG, SST, and UCN3, alongside a notable decrease in CHGA expression in RFX6 KO islets when compared to WT islets (Fig. 6A, B). This indicates that RFX6 is essential for the formation of α-, β-, and δ-cells during development. These reductions were confirmed at the mRNA level, where *INS, GCG, SST,* and *UCN3* were significantly downregulated (Fig. 6C). Furthermore, other key pancreatic islet markers, including *IAPP, PAX6, ARX, GCK, MAFA, KCNJ11, ABCC8, SLC18A1,* and *FEV*, were significantly downregulated (Fig. 6C). On the other, Pancreatic polypeptide Y (*PPY*) was significantly upregulated (Fig. 6C).

**Figure 6.**
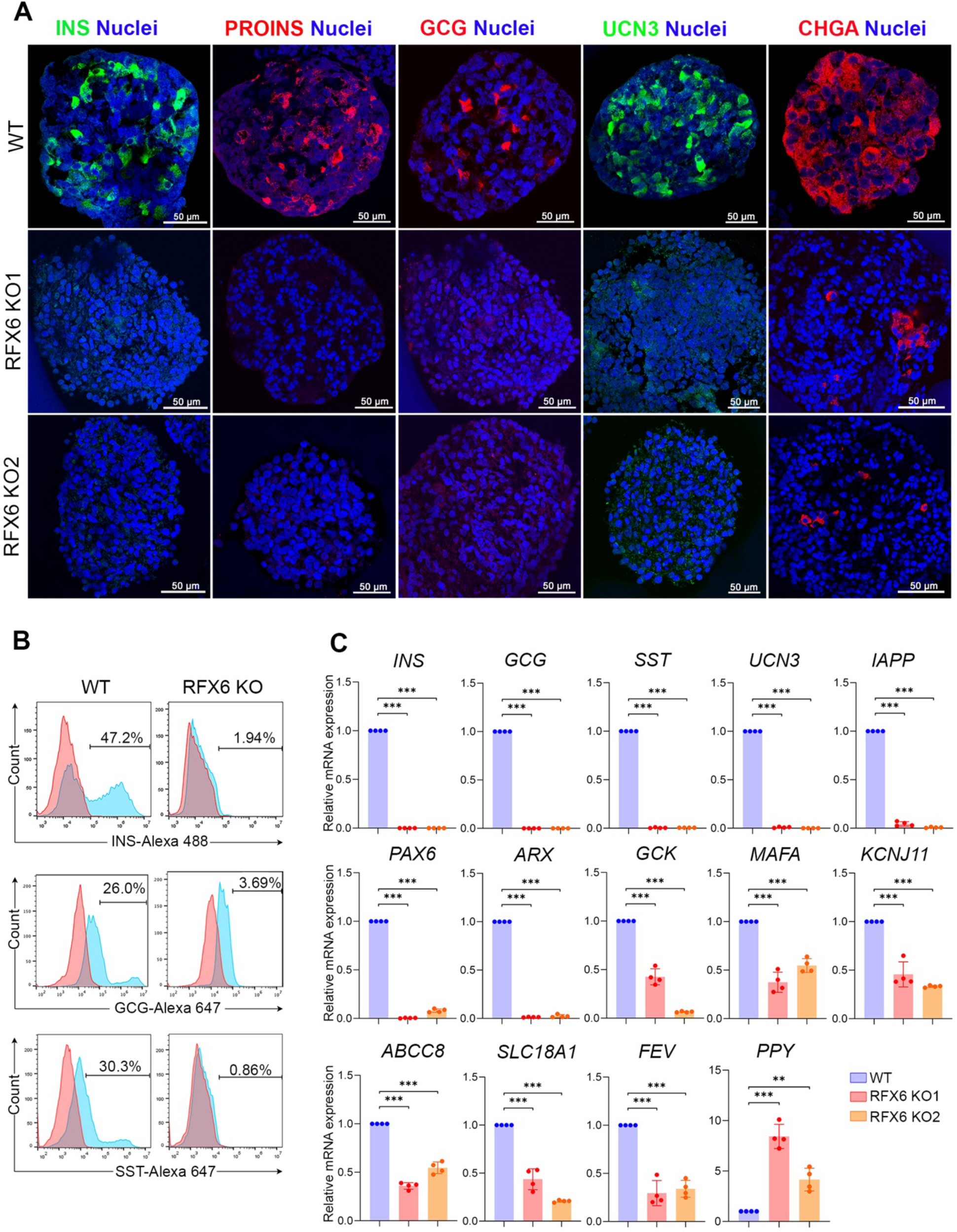
RFX6 loss impairs the development of pancreatic islet cells. **(A)** Confocal immunofluorescence showing expression of pancreatic islet markers, INS, PROINS, GCG, UCN3, and CHGA in islets derived from two different RFX6 KO iPSC lines compared to WT controls (n=3). **(B)** Flow cytometry analysis of the expression of INS, GCG, and SST in islets derived from RFX6 KO iPSCs compared to those derived from WT-iPSCs (n=3). **(C)** RT-qPCR analysis for the mRNA expression of key islet genes, *INS, GCG, SST, UCN3, IAPP, PAX6, ARX, GCK, MAFA, KCNJ11, ABCC8, SCL18A1, FEV,* and *PPY* (n=4). Data are represented as mean ± SD; \**p* < 0.05, \*\**p* < 0.01, \*\*\**p* < 0.001. Scale bar = 100 µm.

### RFX6 overexpression rescues the expression of dysregulated genes in pancreatic progenitors and endocrine progenitors lacking RFX6

We then aimed to assess the impact of ectopic RFX6 expression (RFX6-OE) on reversing the defective phenotypes associated with RFX6 loss in PPs and EPs. RFX6 was overexpressed at day 2 and day 4 of stage 4 of differentiation for assessment of its effect on PPs and EPs, respectively (Fig. 7A). At the end of stage 4, the overexpression of RFX6 resulted in a significant increase in the mRNA expression levels of pancreatic endocrine genes that were downregulated in RFX6 KO PPs, including *RFX6, ARX, PAX6, CHGA, IRX1, IRX2, INS, GCG, SST, MAFB, ERO1B, NEUROD1, PCSK1, ISL1, CRYBA2, SCGN, PTPRN, PTPRN2*, and *LMX1B* (Fig. 7A). Furthermore, we assessed the impact of RFX6-OE on the dysregulated DEGs at day 3 of stage 5, 72 hours post-transfection. Our results revealed a substantial increase in the expression levels of *INS, GCG, SST, NEUROD1, CHGA, CHGB, PAX6, ARX, ISL1, MAFB, PCSK1, ERO1B, IRX2, CRYBA2, KCTD12, LMX1B, SCGN,* and *SSTR2,* following RFX6-OE (Fig. 7B). Moreover, it induced a significant decrease in the *PAX4* mRNA levels, which had been upregulated in RFX6-EPs (Fig. 7B).

**Figure 7.**
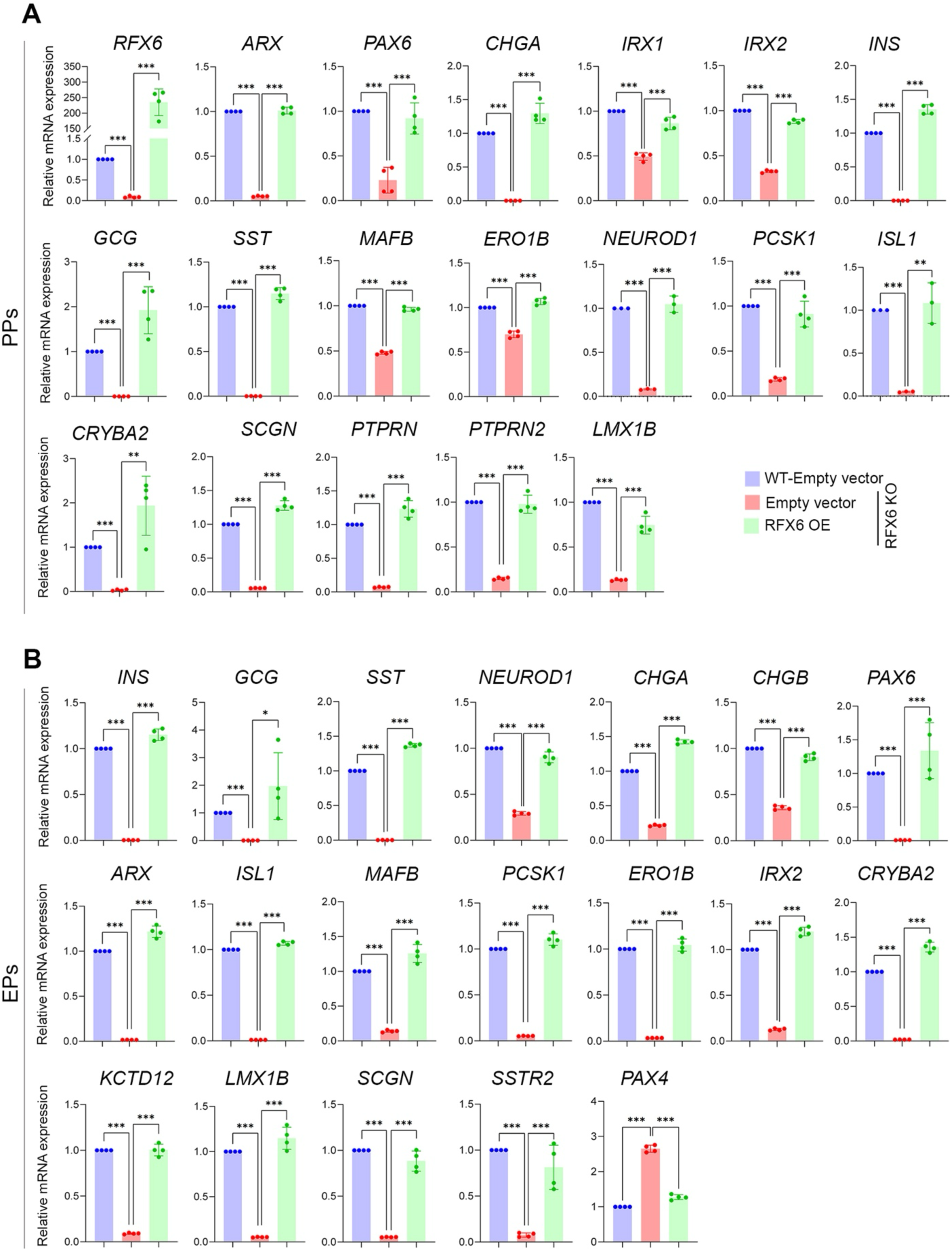
RFX6 overexpression rescues the expression of dysregulated genes in pancreatic cell lacking RFX6. **(A)** RT-qPCR analysis for the expression of pancreatic endocrine genes including, *RFX6*, *ARX*, *PAX6, CHGA, IRX1, IRX2, INS, GCG, SST, MAFB, ERO1B, NEUROD1, PCSK1, ISL1, CRYBA2, SCGN, PTPRN, PTPRN2,* and *LMX1B* in PPs derived from RFX6 KO iPSCs and WT-iPSCs, 48 h following ectopic expression of RFX6 (n=4). **(B)** RT-qPCR analysis for the expression of pancreatic endocrine genes including, *INS, GCG, SST, NEUROD1, CHGA, CHGB, PAX6, ARX, ISL1, MAFB, PCSK1, ERO1B, IRX2, CRYBA2, KCTD12, LMX1B, SCGN, SSTR2,* and *PAX4* in EPs derived from RFX6 KO iPSCs and WT-iPSCs, 72 h following ectopic expression of RFX6 (n=4). Data are represented as mean ± SD; \**p* < 0.05, \*\**p* < 0.01, \*\*\**p* < 0.001.

## Discussion

Recent studies have highlighted the pivotal role of RFX6 in the development and function of human pancreatic islets, linking it to the development of diabetes [24] [25] [17] [3]. Nonetheless, a comprehensive understanding of its specific function during human pancreatic islet development and its exact role in diabetes pathogenesis is still poorly understood. In this study, we precisely examined RFX6 expression across different stages of hPSC-derived pancreatic islets using different approaches. Furthermore, we developed an isogenic KO platform using human iPSC-derived islets to investigate the molecular and cellular alterations at different stages of human islet cell development carrying loss-of-function mutations in RFX6. Our findings are consistent with previous studies, showing the following results: (i) the absence of INS-, GCG-, and SST-producing cells and an increase in PPY cell production due to RFX6 loss; (ii) significant downregulation of genes related to pancreatic endocrine differentiation, insulin secretion, and ion transport in association with RFX6 loss [4, 24, 25]. In addition, our study unveils novel insights into the role of RFX6 during pancreatic islet development. Our data indicate the absence of RFX6 does not impede iPSC differentiation into PPs co-expressing PDX1 and NKX6.1, which serve as precursors to pancreatic β-cells. Furthermore, RFX6 deficiency results in the formation of smaller-sized (hypoplastic) islet organoids, potentially driven by increased cellular apoptosis, likely linked to the deficiency of the antioxidant enzyme CAT. These findings imply that pancreatic hypoplasia and the absence of islet cells due to RFX6 loss-of-function mutations are associated with cellular apoptosis, reduced CAT enzyme expression, and reduced pancreatic endocrine gene expression.

Our findings revealed a significant decrease in PDX1 expression in RFX6 KO-PF compared to WT-PF, consistent with recent findings [25]. However, the absence of RFX6 did not impact the co-expression of PDX1 and NKX6.1 (PDX1^+^/NKX6.1^+^) in PPs. These results align with our timeline expression analysis, which demonstrated the co-localization of RFX6 with PDX1 in the PF stage, while RFX6 showed no co-expression with PDX1 and NKX6.1 in PPs. The difference in the impact on PDX1 expression between PF and PP stages observed in this study may be attributed to RFX6’s involvement during early differentiation stages in intestinal development, as recently reported in iPSC-derived intestinal models [26]. RFX6 is known to be crucial for the development of the small intestine, which shares a common origin with the pancreas, the gut endoderm. In addition to its pancreatic role, PDX1 plays a critical role in small intestine development and function. Previous studies have indicated that during gut tube patterning, PDX1 acts downstream of RFX6, and there is co-expression of PDX1 with RFX6 in the enteroendocrine cells (EECs) of the duodenum and iPSC-derived gut endoderm [27] [28] [26]. Interestingly, RFX6 mutant iPSCs generated defective intestinal organoids due to suppression of PDX1 expression [26]. Our results contradict two prior studies. One such study demonstrated a significant decrease in PDX1 and NKX6.1 levels in PPs derived from MRS patient-specific iPSCs and RFX6 KO-iPSCs [17]. Similarly, another study utilizing RFX6 KO-hESC lines indicated a reduction in the number of PPs due to a marked decrease in PDX1 expression [24]. Our findings suggest that inhibition of PDX1 expression associated with RFX6 loss prior to the PP stage may be linked to defects in intestinal development. This is further supported by the significant reduction in CDX2 expression, which is known for its role in intestinal development within the gut endoderm [29]. Furthermore, these results indicate that RFX6 is not essential for the formation of PDX1+/NKX6.1+ PPs during pancreatic islet development.

The deficiency of RFX6 led to impaired expression of critical TFs and genes essential for endocrine cell development across various stages, including PAX6, INSM1, ARX, NEUROD1, ISL1, IRX1, IRX2, MAFB, TTR, FEV, and CHGA, among others. Conversely, the expression of TFs like PDX1, NKX6.1, SOX9, and FOXA2, specific to PPs [30] remained unaffected by the absence of RFX6 in the PPs, while endocrine TFs like PAX4, NEUROG3, and NKX2.2, were increased in the EPs due to RFX6 deficiency. These findings are consistent with recent results indicating that RFX6 loss does not affect SOX9 expression and increases the expression of NEUROG3, PAX4, and NKX2.2 [25]. During pancreatic development, it has been demonstrated that RFX6 operates downstream of proendocrine TF NEUROG3 [1], which directly controls the expression of PAX4 [31]. Our re-analysis of the previously published single-cell data obtained from different stages of hESC differentiation into pancreatic islets [18] confirmed that the highest expression levels of RFX6 were identified in endocrine clusters, including NEUROG3^high^/GHRL^high^, CHGA^high^/NEUROD1^high^, ERO1LB^high^/ARX^high^, POU2F2^high^/RFX3^high^, TTR^high^/GCG^high^, GCG^high^/CHGA^high^, and HHEX^high^/SST^high^. The analysis also revealed that clusters with high expression levels of PDX1 and SOX9 during the progenitor stages (D11 and D14) did not exhibit RFX6 expression. It has been recently reported that during stage 4 (PPs), a developmental trajectory emerges, leading to the formation of primary endocrine cell groups. The differentiation process becomes notably intricate during stage 5 (EPs), primarily due to the presence of numerous subpopulations [31]. These findings underscore the crucial role of RFX6 in regulating pancreatic endocrine genes important for the development of islet cells, including GCG (α), INS (β), and SST(δ) cells.

Biallelic mutations in RFX6 are associated with permanent neonatal diabetes mellitus (PNDM) in humans, with affected individuals exhibiting smaller size pancreas compared to healthy controls [4]. The cause of this pancreatic hypoplasia is not fully understood. Recent human studies suggested that the reduction in pancreas size may result from the suppression of PDX1 expression at the pancreatic progenitor stage [17, 24]. However, in our current study we demonstrated that biallelic deletion of RFX6 reduced PDX1 in PF, without affecting its expression at PP stage, suggesting the involvement of other mechanisms. Interestingly, our results showed that pancreatic islet organoids derived from RFX6 KO iPSCs was smaller in size compared to WT controls. This size reduction was mainly attributed to increased cell apoptosis during pancreatic endocrine specification, as observed during stage 5 and 6 of differentiation. This result contradicts a previous study used RFX6 KO hESCs, which suggested that the reduced size of the pancreas in patients with RFX6 mutations is not caused by reduced proliferation or increased apoptosis. Instead, it is predominantly attributed to a significant downregulation in PDX1 expression in the early stages of pancreatic development [24]. In our endeavor to unravel the mechanism behind increased cell death in pancreatic cells lacking RFX6, our RNA-seq analysis identified the downregulation of the antioxidant enzyme CAT as a potential cause for the increased cell death. This result was further validated through Western blotting and immunostaining, which revealed almost complete absence of CAT protein in RFX6 KO-PPs and RFX6 KO-EPs compared to WT controls. CAT holds a crucial role in regulating cellular hydrogen peroxide levels, with its enzymatic breakdown acting as a protective mechanism for cells. Particularly, it protects pancreatic β-cells against oxidative damage induced by hydrogen peroxide [32] [33]. Studies have demonstrated that elevated levels of hydrogen peroxide, acting as an oxidant, can inflict harm on pancreatic β-cells and disrupt the signalling pathway involved in insulin production [32, 34]. It has been suggested that the increased levels of hydrogen peroxide associated with mutations in the CAT gene may pose a potential risk for the development of type 2 diabetes, likely due to the peroxide-induced damage to pancreatic β-cells [35]. Taken together, these findings indicate that RFX6 plays a crucial role in safeguarding pancreatic islets during development by maintaining CAT expression, thereby offering protection against oxidative damage.

In summary, our study explored the effects of RFX6 deletion on pancreatic islet development. It showed a substantial reduction in PDX1 expression in RFX6 KO-PF, consistent with earlier studies. However, the absence of RFX6 did not disrupt the development of PDX1 and NKX6.1 in PPs, aligning with the lack of RFX6 co-expression with key progenitor markers and its absence in cell clusters expressing high levels of PDX1 and SOX9 during PPs. The involvement of RFX6 in early intestinal development may shed light on the observed differences in PDX1 expression between PFs and PPs. Furthermore, our findings indicate that RFX6 regulates the expression of crucial pancreatic endocrine genes essential for the formation of INS-, GCG-, and SST-expressing cells during pancreatic differentiation. Moreover, RFX6 deletion resulted in smaller pancreatic islet organoids, attributed to increased cell apoptosis during endocrine specification. This apoptotic process was associated with a significant reduction in CAT expression, which protects pancreatic cells from oxidative damage. These results underscore RFX6’s pivotal role in safeguarding pancreatic islets, potentially explaining pancreatic hypoplasia in patients with RFX6 homozygous mutations. Thus, our study highlights the complexity of RFX6’s role in pancreatic islet development and its implications for understanding pancreatic hypoplasia and diabetes risk.

## Methods

### Culture and maintenance of hESCs and iPSCs

HA-RFX6 tagged H9 hESC line (RFX6^HA/HA^ H9-hESCs) with its control, H9-hESCs, were obtained from Dr. N. Ray Dunn (A*STAR, Singapore). Wild type iPSCs (WT-iPSCs) generated and fully characterized in our laboratory were used [21, 36]. Cells were cultured and maintained in mTeSR Plus medium (Stem Cell Technologies, Canada) on Matrigel-coated dishes (Corning, USA), following our established approach [37][38].

### Generation of RFX6 knockout iPSCs

For RFX6 knockout generation, we generated two RFX6 KO hiPSC lines (RFX6 KO1 and RFX6 KO2) from WT-iPSCs. Cells were transfected with a GFP-tagged plasmid vector expressing spCas9 and gRNA using Lipofectamine 3000 Transfection Reagent following manufacturer’s protocol (ThermoFisher Scientific). Cells were then sorted based on GFP expression after 48 hours from the transfection.

### Differentiation of hESCs and iPSCs into pancreatic islets

hESC/iPSC lines were differentiated *in vitro* into PPs using our established protocol [39]. Veres et al. protocol was adapted for further differentiation into pancreatic islet cells [40]. iPSCs were differentiated in a 2D culture system until the end of PP stage. Later, cells were dissociated using TryplE into single cells, counted, and organoids were formed using Aggrewell 400 24-well plates (Stem Cell Technologies). After counting, 2×10^6^ cells were divided into each well of 24-well plate, ∼1,666 cells per microwell. Cells were kept in the aggrewell plate for 48 hours then transferred into an ultra-low attachment plate where differentiation was continued. Cells were incubated on a shaker after organoid formation. The list of differentiation reagents used are listed in Table S1, Supporting information.

### Paraffin embedding and immunofluorescence

For cultured cells, immunostaining was conducted on cells that had been fixed using 4% paraformaldehyde (PFA) in PBS for 20-25 min, following a protocol described earlier [37]. Briefly, cells were permeabilized with PBS containing 0.5% Triton (PBST) for 15 minutes and were blocked overnight at 4°C using 6% bovine serum albumin (BSA) in PBST. Primary antibodies were added overnight at 4°C then cells were washed three times using TBST containing 0.5% Tween (TBST). Secondary antibodies were added at room temperature for 1 hour followed by three washes with TBST. Nuclear staining was done using Hoechst 33258 diluted 1:5000 in PBS (Life Technologies, USA). Cells were washed using PBS and imaged using inverted fluorescence microscope (Olympus). Used antibodies’ details are listed in Table S2, Supporting information.

For the 3D pancreatic islet organoids, we employed the paraffin embedding technique. Islet organoids were collected into a 1.5 mL Eppendorf tube, washed with PBS, and subsequently fixed with 4% PFA for 20-30 minutes on a shaker at room temperature. After fixation, the organoids underwent two washes with TBST followed by the addition of PBS The organoids were then mixed with 100 µL of melted Histogel (Epredia) in a 7×7×5 mm disposable embedding mold. These histogel blocks were released from the mold and wrapped in a 3-ply tissue and inserted into tissue cassettes. The dehydration series for the blocks was adapted from Kong et al.’s paraffin embedding protocol [41]. The paraffin blocks were then sectioned using a microtome at a thickness of 5 µm. Antigen retrieval was performed using the method adapted from Campbell-Thompson et al.’s [42]. Subsequently, slides were permeabilized using PBST for 15 minutes at room temperature and blocked with 6% BSA in PBST for at least 2 hours at room temperature. The subsequent steps followed were similar to those in immunofluorescence.

### Flow cytometry

Differentiated cells were collected at different stages from one well of a 6-well plate. Cells were first washed once with PBS and then dissociated using TrypLE into single cells. Afterwards, cells were collected in a 15 mL Falcon Tube (ThermoFisher Scientific, USA) and resuspended in 1 mL of PBS. All cell centrifugations were done at 1800 rpm for 5 minutes (Eppendorf). Cells were resuspended in 200 µL of cold PBS and then 1 mL of 4% PFA was added then incubated, slanted, at room temperature for 20 minutes on a shaker. Cells were washed twice with TBST then permeabilized using PBST. Cells were blocked using 6% BSA in PBST overnight at 4°C. 1-2×10^5^ of cells, around 100 µL, were transferred into V-bottom shaped 96-well plate (Corning, USA) and primary antibodies were added for 3 hours on a shaker, at room temperature. Cells were washed twice with TBST then incubated with secondary antibodies diluted in PBS for 30 minutes at room temperature. Cells were washed again twice and resuspended in PBS then run using BD Accuri C6 Flow Cytometer then analysed using FlowJo software. Used antibodies’ details are listed in Supplementary Table 2.

### Western blotting

Total protein was extracted from 1-2 wells of a 6-well plate using RIPA lysis buffer with protease inhibitor (ThermoFisher Scientific). Protein concentration was measured using Pierce BCA kit (ThermoFisher Scientific, #23225). 20-30 µg of total protein were loaded and separated using 7.5-10% SDS-PAGE gels then transferred onto PVDF membranes (ThermoFisher Scientific, #88518). Membranes were blocked using 15% skimmed milk in TBST at least for 3 hours at room temperature or overnight at 4°C. Primary antibody was then added and incubated overnight at 4°C. Membranes were then washed using TBST and secondary antibody was added for 1 hour at room temperature followed by more TBST washes. Membranes were developed using SuperSignal West Pico Chemiluminescent substrate (ThermoFisher Scientific, #34580). Used antibodies’ details are listed in Supplementary Table 2.

### RNA extraction, PCR, and RT-qPCR

Cells were collected from one well of a 6-well plate using 700 µL of TRIzol Reagent (Life Technologies) and RNA was extracted using Direct-zol RNA Miniprep (Zymo Research). 1µg of total RNA was used for cDNA synthesis using High-Capacity cDNA Reverse Transcription Kit while following manufacturer’s protocol (Applied Biosystems). PCR was done using 2X PCR Master Mix (Thermofisher Scientific, #K0171). RT-qPCR was performed using GoTaq qPCR SYBR Green Master Mix (Promega). GAPDH was used as an endogenous control (primer details are listed in Supplementary Table 3).

### RNA-sequencing analysis

The mRNA isolation utilized the NEBNext Poly(A) mRNA Magnetic Isolation Kit (NEB, #E7490) with 1 μg of total RNA. Subsequently, RNA-seq libraries were generated using the NEBNext Ultra Directional RNA Library Prep Kit (NEB, #E7420L), followed by sequencing on an Illumina Hiseq 4000 system. Raw data underwent conversion to FASTQ files through Illumina BCL2Fastq Conversion Software v2.20. For intial preprocessing of the Pair-end FASTQ files, nf-core/rnaseq (version 2.7.2) pipeline implemented in the nextflow workflow (version 23.10.1) was used. STAR (version 2.7.9a) was used for the read alignments, Salmon (version 1.5.0) for quantification of reads, TrimGalore (version 0.6.6) was applied for read trimming and GENCODE (version 38) was used for annotation of genes [43]. The count matrix generated by Salmon was filtered, excluding genes labeled ’Mt_tRNA,’ Mt_rRNA,’rRNA’ and ’rRNA_pseudogene’ in the GENCODE annotation file. Subsequently, low expression genes were excluded using the HTSFilter (version 1.32.0) [44]. Differential expression analysis were performed using the DESeq2 (version 1.32.0) to identify the DEGs for specific comparisons. PCA was generated using the glmpca package of R. DEGs were identified based on criteria of log2 fold change (FC) > 1 and <-1, with a P-value < 0.05. Subsequent analyses also included Gene Ontology (GO) and Kyoto Encyclopaedia of Genes and Genomes (KEGG) pathways, performed using the Database for Annotation, Visualization, and Integrated Discovery (DAVID) [45].

### Apoptosis and proliferation assays

The apoptosis assay was performed using the Annexin V-FITC Apoptosis Detection Kit (Abcam, #ab14085), while the proliferation assay was conducted employing the BrdU incorporation method (Invitrogen, #000103), following previously established protocols [46]. For the proliferation assay, cells were exposed to 20 µM BrdU in differentiation media for 5-6 hours at 37°C. Subsequently, the cells were washed with PBS, dissociated using TryplE, and fixed with cold 70% ethanol overnight at 4°C. Following this, the cells were rinsed once with PBS and denatured with 0.2 M HCL containing 0.5% Triton for 15-20 minutes at room temperature, followed by a 0.1 M sodium tetraborate treatment for another 15-20 minutes. Afterward, the cells were washed with PBS and blocked using 6% BSA in PBS with 0.1% saponin. Cells were then incubated with Alexa Fluor 488-conjugated BrdU monoclonal antibody (Thermo Fisher, #B35130, 1:100) overnight at 4°C. Cells were then washed with TBST twice and quantification of BrdU-positive cells was conducted through flow cytometry analysis using BD Accuri C6 Flow Cytometer.

### Overexpression of RFX6

Cells were trypsinized and resuspended in differentiation media with 10 µM Y-27632. Resuspended cells were transfected with either RFX6 plasmid (RFX6 (Myc-DDK-tagged), RC206174, OriGene, USA) or empty vector. Lipofectamine 3000 Transfection Reagent was used while following manufacturer’s protocol (Thermofisher Scientific, #L3000-015). For stage 4 transfection, cells were transfected at the end of day 2 of stage 4 and collected after 48 hours (day 4 of stage 4), while in stage 5 transfection, cells were transfected at end of stage 4 and collected after 72 hours (day 3 of stage 5).

### Single cell analysis

The online published GSE202497 data set (https://www.ncbi.nlm.nih.gov/geo/query/acc.cgi?acc=GSE202497) was used [18] and re-analyzed. In total, 25686 cells were retrieved from the dataset. The data consists of 5 samples from human pluripotent stem cell differentiation into islets at D11, D14, D21, D32, D39 days. Top 2000 highly variable genes were obtained using Seurat V3 algorithm implement in Scanpy. The data was log-normalized and Harmony with “sample” as batch variable was used for technical batch-effect correction. Neighborhood graph was computed using the sc.pp.neighbors function using top 50 batch-effect adjusted principal components. Clustering at different resolutions was performed using the leiden algorithm. Marker genes for each cluster were calculated using the Wilcoxon method implemented in rank_genes_groups function. Finally, using the marker genes, the clusters were manually annotated. Top marker genes and cluster names are provided in Supplementary Figure 2.

### Analysis of protein-protein interaction networks associated with the CAT gene

To explore potential interactions and predict functional associations related to the CAT gene, we employed the STRING database (https://string-db.org) [23].

### Statistical analysis

At least three biological replicates were used in most experiments while statistical analysis was done using unpaired two-tailed Student’s t-test on Prism 8. Data are represented as the mean ± standard deviation (SD).

## Supporting information

Suppelemtary Figures and Tables

## Acknowledgements

We would like to thank Dr. N. Ray Dunn (A*STAR, Singapore) for providing the HA-RFX6 tagged H9 hESC lines (RFX6^HA/HA^ H9-hESCs). Furthermore, we thank the Genomic Core members at QBRI for their assistance for technical support in RNA sequencing.

## Data Availability

RNA-seq datasets have been deposited in the Zenodo repository with accession link (DOI: 10.5281/zenodo.10656891).

## Authors’ relationships and activities

SH is a co-founder and shareholder of Sequantrix GmbH and has research funding from by Novo Nordisk and Askbio. The authors declare that there are no relationships or activities that might bias, or be perceived to bias, their work.

## Contributions statement

N. A. performed most of the experiments, analyzed the data. A.K.E. and B.M. performed experiments and analyzed the data. S.I. and S.H. analyzed the sequencing data. E.M.A. Conceived and designed the study, supervised the project, analyzed and interpreted the data, and wrote the manuscript. All authors approved the final version of the manuscript.

