## Supplementary material for "Deletion of RFX6, a Diabetes-Associated Gene, Impairs iPSC-Derived Islet Organoid Development and Survival, With No Impact on the Generation of PDX1+/NKX6.1+ Progenitors": Suppelemtary Figures and Tables

### Supplementary Figures

#### Supplementary Figure 1

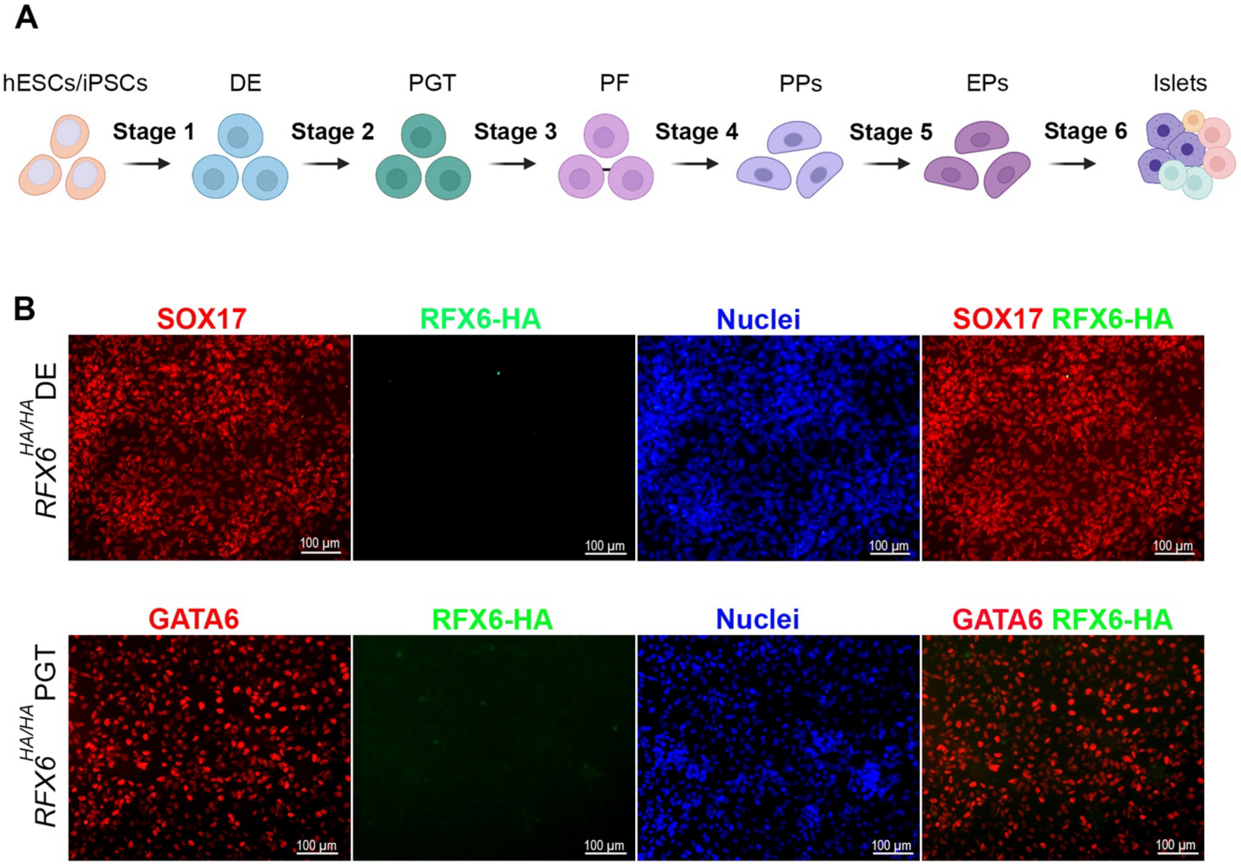


**Supplementary Figure 1**. Expression of RFX6 during early stages of hESC differentiation. (A) Diagram illustrating the procedural outline for the differentiation of iPSCs/hESCs into pancreatic islets. (B) Immunostaining showing the expression of RFX6 during differentiation of hESC-H9 into stage 1 (DE) and stage 2 (PGT). DE: definitive endoderm, PGT: primitive gut tube, PF: posterior foregut, PPs: pancreatic progenitors, EPs: endocrine progenitors. Scale bars = 100 µm.

**Supplementary Figure 2**

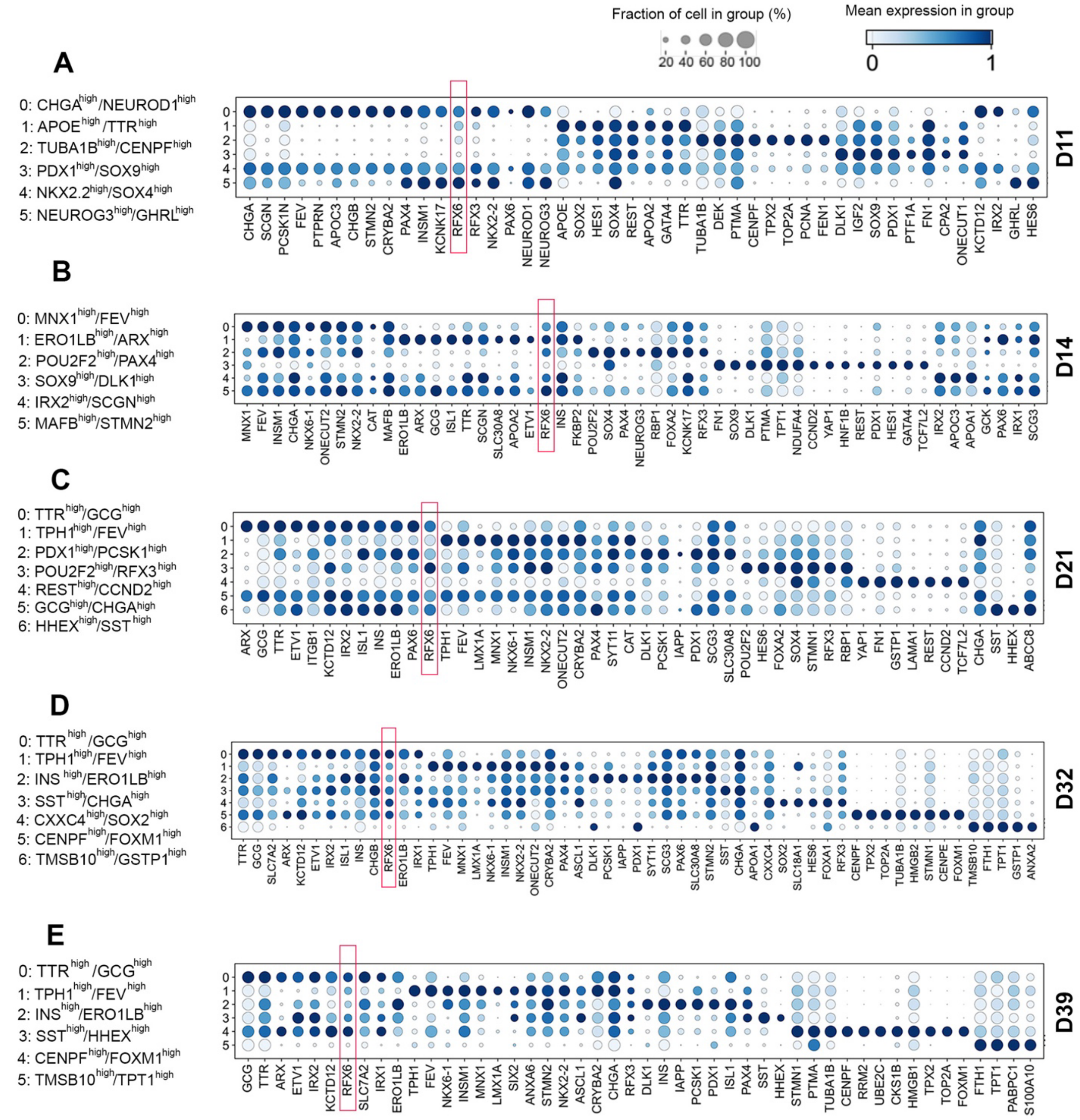


**Supplementary Figure 2.** Single-cell sequencing analysis of hESC-derived pancreatic islets. Dotplot showing marker gene distributions across the different cell populations at different days of differentiation, including day 11 (D11) (A), day 14 (D14) (B), day 21 (D21) (C), day 32 (D32) (D), and day 39 (D39) (E).

**Supplementary Figure 3**

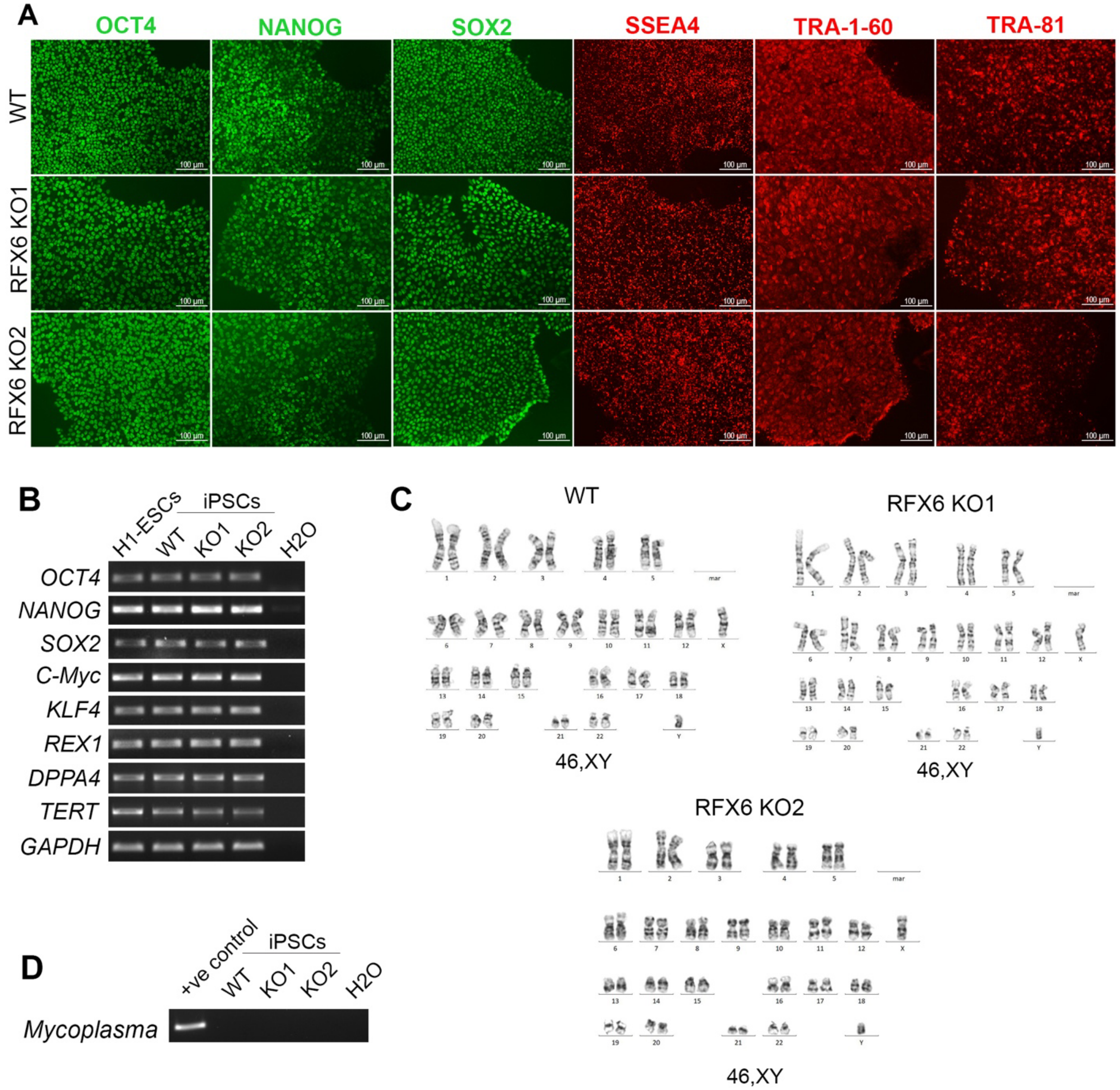


**Supplementary Figure 3.** CRISPR/Cas9-edited iPSC lines are pluripotent. (A) Immunofluorescence staining showcases the expression of pluripotency markers OCT4, NANOG, SOX2, SSEA4, and TRA-1-81 in both WT and RFX6 KO iPSC clones, with nuclei counterstained using Hoechst (blue). (B) RT-PCR analysis depicts the expression of pluripotency-associated genes OCT4, NANOG, SOX2, C-MYC, KLF4, REX1, DPPA4, and TERT. (C) Karyotype analysis reveals a normal karyotype in both WT and RFX6 KO iPSC clones. (D) PCR analysis confirms the absence of mycoplasma contamination in the WT and RFX6 KO iPSC lines. Scale bar = 100 μm.

**Supplementary Figure 4**

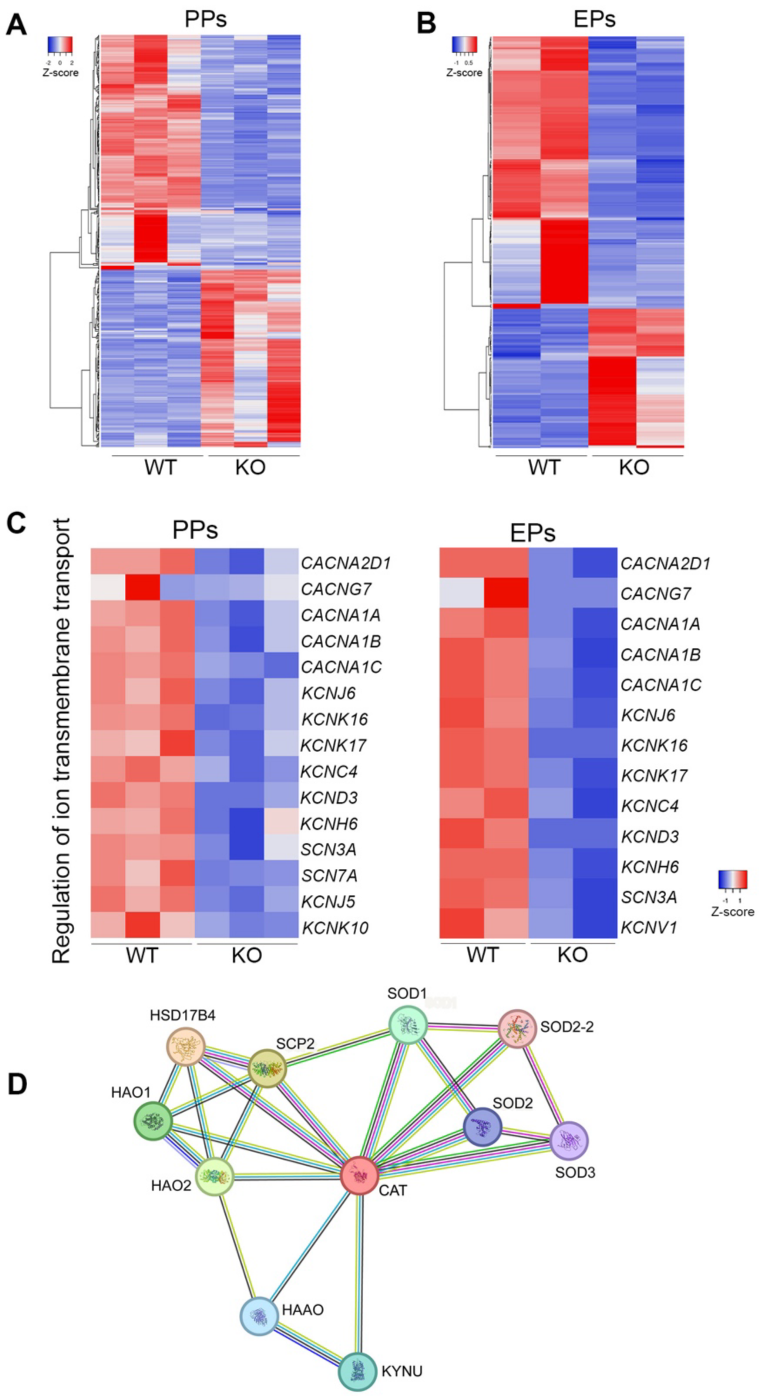


**Supplementary Figure 4.** RNA-seq analysis of pancreatic progenitors (PPs) and endocrine progenitors (EPs) derived from WT and RFX6 KO iPSCs. A clustering heatmap of differentially expressed genes (DEGs) in PPs (A) and EPs (B) derived from WT and RFX6 KO iPSCs. (C) Heatmaps of DEGs associated with the regulation of ion transmembrane transport in RFX6 KO PPs and RFX6 KO-EPs compared to WT controls (p-value <0.05 and Log2 FC< -1.0). (D) STRING network analysis showing the interaction of Catalase (CAT) protein with other proteins.

**Supplementary Table 1.** *In vitro* pancreatic differentiation protocol.

| Stage of Differentiation | Media | Final Cytokine Concentration |
|---|---|---|
| Stage 1:<br>Definitive Endoderm<br>(Day 1) | MCDB131<br>1% Pen/Strep<br>1% L-Glutamine<br>10 mM Glucose<br>0.5% BSA<br>1.5 g/L $NaHCO_3$ | 100 ng/mL Activin A<br>0.25 mM Vitamin C<br>2 µM CHIR99021<br>1 mM Rock Inhibitor |
| Stage 1:<br>Definitive Endoderm<br>(Days 2-3) | MCDB131<br>1% Pen/Strep<br>1% L-Glutamine<br>10 mM Glucose<br>0.5% BSA<br>1.5 g/L $NaHCO_3$ | 100 ng/mL Activin A<br>0.25 mM Vitamin C |
| Stage 2:<br>Primitive Gut Tube<br>(Days 4-5) | MCDB131<br>1% Pen/Strep<br>1% L-Glutamine<br>10 mM Glucose<br>0.5% BSA<br>1.5 g/L $NaHCO_3$ | 3 ng/mL Wnt3a<br>0.75 µM Dorsomorphin<br>0.25 mM Vitamin C<br>50 ng/mL FGF10 |
| Stage 3:<br>Posterior Foregut<br>(Days 6-7) | DMEM<br>1% Pen/Strep<br>1% L-Glutamine<br>4.5 g/L D-Glucose<br>110 mg/L Sodium Pyruvate | 1% B27<br>200 nM LDN<br>0.25 mM Vitamin C<br>50 ng/mL FGF10<br>2 µM Retinoic acid<br>0.25 µM SANT-1 |
| Stage 4:<br>Pancreatic Progenitor<br>(Days 8-11) | DMEM<br>1% Pen/Strep<br>1% L-Glutamine<br>4.5 g/L D-Glucose<br>110 mg/L Sodium Pyruvate | 1% B27<br>200 nM LDN<br>100 ng/mL EGF<br>0.25 mM Vitamin C<br>10 mM Nicotinamide |
| Stage 5:<br>Endocrine<br>Progenitor<br>(Days 12-15) | MCDB131<br>1% Pen/Strep<br>1% L-Glutamine<br>20% BSA<br>1.754 g/L $NaHCO_3$<br>20 mM Glucose<br>5 mL ITS<br>10 mg/L Heparin | 0.25 µM SANT-1<br>20 ng/mL Beta-Cellulin<br>1 µM Gamma secretase inhibitor<br>10 µM Alk5i<br>1 µM T3<br>0.1 µM Retinoic acid<br>0.25 mM Vitamin C |
| Stage 5: | MCDB131 | 20 ng/mL Beta-Cellulin |

| | | |
|---|---|---|
| Endocrine Progenitor (Days 16-18) | 1% Pen/Strep<br>1% L-Glutamine<br>20% BSA<br>1.754 g/L $NaHCO_3$<br>20 mM Glucose<br>5 mL ITS<br>10 mg/L Heparin | 1 μM Gamma secretase inhibitor<br>10 μM Alk5i<br>1 μM T3<br>0.1 μM Retinoic acid<br>0.25 mM Vitamin C |
| Stage 6: Pancreatic Islet (Days 19-32) | MCDB131<br>1% Pen/Strep<br>1% L-Glutamine<br>20% BSA<br>1.23 g/L $NaHCO_3$<br>2.5 mM Glucose<br>5 mL ITS | 0.25 mM Vitamin C |

**Supplementary Table 2.** List of used antibodies for immunostaining and western blot.

| **Antibody** | **Company** | **Catalog No.** | **Dilution** |
|---|---|---|---|
| B-actin | Santa Cruz | Sc-47778 | WB (1:10,000) |
| CAT | Cell Signaling | 12980S | WB (1:4000)<br>IS (1:1000) |
| CDX2 | Abcam | ab76541 | WB (1:2500)<br>IS (1:1000) |
| CHGA | ThermoFisher | MA5-14536 | WB (1:4000)<br>IS (1:3000) |
| FOXA2 | Cell Signaling | 3143 | WB (1:4000)<br>IS (1:1000) |
| FOXA2 | Abcam | ab60721 | IS (1:1000) |
| GATA6 | R&D | AF1700 | IS (1:500) |
| GCG | Sigma | G2654 | IS (1:2000)<br>FACs (1:100) |
| HA | Cell Signaling | 3724s | IS (1:1000) |
| INS | DSHB | GN-ID4-s | IS (1:2000)<br>FACs (1:100) |
| OCT4 | Cell Signaling | 9656 | IS (1:500) |
| NANOG | Cell Signaling | 9656 | IS (1:500) |
| NGN3 | R&D | AF3444 | IS (1:1000) |
| NKX2.2 | DSHB | 74.5A5-c | IS (1:2000) |
| NKX6.1 | DSHB | F55A12-C | WB (1:4000)<br>IS (1:2000)<br>FACs (1:100) |
| PDX1 | Abcam | ab47308 | WB (1:4000) / (1:1000)<br>IS (1:500)<br>FACs (1:100) |
| PROINS | R&D | MAB13361 | IS (1:2000) |
| RFX6 | Sigma | HPA037696 | WB (1:200) |

| | | | |
|---|---|---|---|
| SOX2 | Cell Signaling | 9656 | IS (1:500) |
| SOX9 | Sigma | HPA001758 | WB (1:4000) |
| SOX17 | Origene | CF500096 | IS (1:500) |
| SSEA4 | Cell Signaling | 9656 | IS (1:500) |
| SST | Millipore | MAB354 | FACs (1:100) |
| TRA-1-60 | Cell Signaling | 9656 | IS (1:500) |
| TRA-81 | Cell Signaling | 9656 | IS (1:500) |
| UCN3 | Sigma | HPA038281 | IS (1:1000) |
| 594 donkey anti-rabbit | Invitrogen | A21207 | IS (1:500) |
| 488 donkey anti-rabbit | Invitrogen | A21206 | IS (1:500) |
| 568 donkey anti-mouse | Invitrogen | A10037 | IS (1:500) |
| 488 donkey anti-mouse | Invitrogen | A21202 | IS (1:500)<br>FACs (1:500) |
| 647 donkey anti-mouse | Invitrogen | A31571 | FACs (1:500) |
| 568 donkey anti-goat | Invitrogen | A11057 | IS (1:500) |
| 488 donkey anti-sheep | Invitrogen | A11015 | IS (1:500) |
| 488 goat anti-rat | Invitrogen | A11006 | IS (1:500)<br>FACs (1:500) |
| 568 goat anti-rat | Invitrogen | A11077 | IS (1:500) |
| 647 donkey anti-rat | Invitrogen | A48272 | FACs (1:500) |
| 488 goat anti-guinea pig | Invitrogen | A11073 | IS (1:500) |

| | | | |
|---|---|---|---|
| 647 goat anti-guinea pig | Invitrogen | A21450 | FACs (1:500) |
| Peroxidase AffiniPure Donkey anti-Rabbit IgG (H+L) | Jackson ImmunoResearch Laboratories | 711-035-152 | WB (1:10,000) |
| Peroxidase AffiniPure Donkey anti-Mouse IgG (H+L) | Jackson ImmunoResearch Laboratories | 715-035-150 | WB (1:10,000) |
| Peroxidase AffiniPure Donkey anti-guinea pig IgG (H+L) | Jackson ImmunoResearch Laboratories | 706-035-148 | WB (1:10,000) |

**Supplementary Table 3.** List of primers used for RT-qPCR validation.

| Gene | Forward | Reverse |
|---|---|---|
| *ABCC8* | TTGCCGAAACCGTAGAAGG | CGGTTCCAGCAGAAGCTTC |
| *ARX* | CTGCTGAAACGCAAACAGAGGC | CTCGGTCAAGTCCAGCCTCATG |
| *CDX2* | CTGGAGCTGGAGAAGGAGTTTC | ATTTTAACCTGCCTCTCAGAGAGC |
| *CHGA* | GAAGAAGGCCCCACTGTAGT | TTCCCAGCTCCATCCACAG |
| *CHGB* | CACGCCATTCTGAGAAGAGC | TCTCCTGGCTCTTCAAGGTG |
| *CPA1* | ACTACGCCACCTACCACACC | GGTGTTGCCAATCTGGATCT |
| *CPA2* | ATCTTCCTCCTGCCAGTCAC | CACACCAACACAGAGGCTTC |
| *CRYBA2* | GATGTGGGTTCCCTCAAAGT | GCTCACCGTAAGTACAGAACTC |
| *ERO1B* | TGAACCCAGAGCGTTACACT | GGCGCCAGAGGATTTAAAGG |
| *FEV* | GCCTCTCCAAACTCAACCTC | CAAGCTGGGACTGGGGTAG |
| *FOXA2* | GGGAGCGGTGAAGATGGA | TCATGTTGCTCACGGAGGAGTA |
| *GAPDH* | ACGACCACTTTGTCAAGCTCATTTC | GCAGTGAGGGTCTCTCTCTTCCTCT |
| *GCG* | CTCTTCACCTGCTCTGTTCTAC | TGGATTTCTCCTCTGTGTCTTG |
| *GCK* | GCATCTTCCAGCTCTTCGAC | GGGCTACATTTGAAGGCAGA |
| *IAPP* | TTGAGAAGCAATGGGCATCC | GGGTGTAGCTTTCAGATGGTTC |
| *INS* | AAGAGGCCATCAAGCAGATCA | CAGGAGGCGCATCCACA |
| *INSM1* | TTTGTCTCGTGGTTGGAAGC | CCAAAACAACCCGTACGCTA |
| *IRX1* | CAAGAATCCCTACCCCACCA | TCCCCATGTCACCTTGTTCT |
| *IRX2* | TCACCAAGATGACCCTCACC | TTCGCTTTTGTTTCTCGGGG |
| *ISL1* | CTGTGGACATTACTCCCTCTTAC | GCAACCAACACATAGGGAAATC |
| *KCNJ11* | GCGCTTTGTGCCCATTGTA | TTGACGGTGTTGCCAAACTTG |
| *KCTD12* | GCCTCTGTCACTCAAGTCTTAC | TGACAGGGTGCAGTGAATAAG |
| *LMX1B* | ACCTCCTTAACCAGCCTCAG | GCATGGAGTAGAGCCGGTC |
| *MAFA* | GCACATTCTGGAGAGCGAGA | CCGCCAGCTTCTCGTATTTC |
| *MAFB* | GGAGAATGAGAAGACGCAGC | GTTTCTCGCACTTGACCTTGT |
| *NEUROD1* | GCCCCAGGGTTATGAGACTAT | GAGAACTGAGACACTCGTCTGT |
| *NEUROG3* | GGCTGTGGGTGCTAAGGGTAAG | CAGGGAGAAGCAGAAGGAACAA |
| *NKX2.2* | AAACCATGTCACGCGCTCA | GGCGTTGTACTGCATGTGCT |
| *NKX6.1* | GGGCTCGTTTGGCCTATTCGTT | CCACTTGGTCCGGCGGTTCT |
| *ONECUT2* | GCCATCTTCAAGGAGAACAAAC | CGTTCATGAAGAAGTTGCTGAC |

| | | |
|---|---|---|
| *PAX4* | AGCAGAGGCACTGGAGAAAGAGTT | CAGCTGCATTTCCCACTTGAGCTT |
| *PAX6* | GCGGAAGCTGCAAAGAAATAG | GGGCAAACACATCTGGATAATG |
| *PCSK1* | TCACACATGGGGAGAGAACC | TCCCGTGCAAAATCAGCTTC |
| *PDX1* | CGTCCAGCTGCCTTTCCCAT | CCGTGAGATGTACTTGTTGAATAGGA |
| *PPY* | AGGTGCTCGCTTGGTCTAGTG | ACCCAGCAGTGGCTGTAGTAAC |
| *PTPRN* | CCTACCAAGCAGAGCCAAAC | TGGTCATAGGGCAGGAAGTC |
| *PTPRN2* | ACTGAGGATGTGGAGAAGGC | TGAGTTTGCTTTTCGACCCG |
| *RFX6* | GTCGATGCATGGCTTGGACT | TGGGCCATAGCTAGACGGTG |
| *SCGN* | CTGGGTACTGATGACACGGT | CCAGCAAGCTCTTTCATCCG |
| *SIX3* | GCGACTCGGAATGTGATGTAT | GGAGAAGGAAGAGGAGGAAGA |
| *SLC18A1* | ATGGTCATCACTGGGGTCAT | GGCTTCTGGGTTGCATACAT |
| *SOX2* | GGGAAATGGGAGGGGTGCAAAAGAGG | TTGCGTGAGTGTGGATGGGATTGGTG |
| *SOX9* | GACTACACCGACCACCAGAACTCC | GTCTGCGGGATGGAAGGGA |
| *SST* | AGCTGCTGTCTGAACCCAAC | CCATAGCCGGGTTTGAGTTA |
| *SSTR2* | GCTGTGCCAACCCTATCCTA | TCCTGCTTACTGTCACTCCG |
| *TTR* | CATGGGCTCACAACTGAGGA | TTGGCTGTGAATACCACCTCTG |
| *UCN3* | GATGGGCTTGGCTTTGTAGA | GGAGGGAAGTCCACTCTCG |
